# A whole-genome scan for evidence of recent positive and balancing selection in aye-ayes (*Daubentonia madagascariensis*) utilizing a well-fit evolutionary baseline model

**DOI:** 10.1101/2024.11.08.622667

**Authors:** Vivak Soni, John W. Terbot, Cyril J. Versoza, Susanne P. Pfeifer, Jeffrey D. Jensen

**Affiliations:** Center for Evolution and Medicine, School of Life Sciences, Arizona State University, Tempe, AZ, USA

**Keywords:** primate, aye-aye, selective sweep, balancing selection, genome scan, demography

## Abstract

The aye-aye (*Daubentonia madagascariensis*) is one of the 25 most endangered primate species in the world, maintaining amongst the lowest genetic diversity of any primate measured to date. Characterizing patterns of genetic variation within aye-aye populations, and the relative influences of neutral and selective processes in shaping that variation, is thus important for future conservation efforts. In this study, we performed the first whole-genome scans for recent positive and balancing selection in the species, utilizing high-coverage population genomic data from newly sequenced individuals. We generated null thresholds for our genomic scans by creating an evolutionarily appropriate baseline model that incorporates the demographic history of this aye-aye population, and identified a small number of candidate genes. Most notably, a suite of genes involved in olfaction — a key trait in these nocturnal primates — were identified as experiencing long-term balancing selection. We also conducted analyses to quantify the expected statistical power to detect positive and balancing selection in this population using site frequency spectrum-based inference methods, once accounting for the potentially confounding contributions of population history, recombination and mutation rate variation, and purifying and background selection. This work, presenting the first high-quality, genome-wide polymorphism data across the functional regions of the aye-aye genome, thus provides important insights into the landscape of episodic selective forces in this highly endangered species.

## INTRODUCTION

A strepsirrhine endemic to Madagascar and the world’s largest nocturnal primate, the aye-aye (*Daubentonia madagascariensis*) is the only extant member of the *Daubentoniidae* family, exhibiting a geographical range wider than any other member of the Lemuroidea superfamily (Sterling 1993). However, rapid habitat destruction is thought to have contributed to a severe population decline over the past few decades (Louis et al. 2020; Suzzi-Simmons 2023), along with direct human predation at least partially owing to the regional Malagasy cultural belief that aye-ayes are a harbinger of illness and death (Andriamasimanana 1994). These ongoing trends – coupled with harboring amongst the lowest genetic diversity of any primate measured to date (Perry et al. 2012; Kuderna et al. 2023) – have placed the aye-aye on the list of the 25 most endangered primate species in the world, according to the International Union for Conservation of Nature and Natural Resources Species Survival Commission Primate Specialist Group (Schwitzer et al. 2013; Louis et al. 2020; and see the discussion in Gross 2017). As such, characterizing patterns of genetic variation within aye-aye populations, and the relative influences of neutral and selective processes in shaping that variation, will be important for future conservation efforts.

### Signatures of episodic selection

Although it is well understood that the demographic history of a population together with the recurrent action of natural selection acts to shape patterns of polymorphism at the DNA sequence level, disentangling the effects of these processes remains an ongoing concern (e.g., Ewing and Jensen 2016; Charlesworth and Jensen 2022; Jensen 2023). Nevertheless, distinguishing these processes is fundamental for gaining an improved understanding of general evolutionary dynamics, inasmuch as it would improve our understanding of the relative importance of adaptive and nonadaptive factors in shaping levels of genetic variation in natural aye-aye populations as well as facilitate the accurate identification of genomic regions that have experienced recent bouts of episodic selection. Existing methods for detecting recent beneficial fixations rely on the changes in patterns of variation at linked sites (i.e., by characterizing the resulting effects of the associated selective sweep), though the nature of these changes will naturally depend on the details of the selective pressure (see the review of Stephan 2019). The term "selective sweep" describes the process whereby a positively selected mutation rapidly increases in frequency and fixes within a population, with linked variation following the same trajectory to an extent determined by the level of linkage (Maynard Smith and Haigh 1974; and see the review of Charlesworth and Jensen 2021). The fixation of these linked variants is expected to temporally reduce local nucleotide diversity (Berry et al. 1991). Under a single selective sweep model with recombination, there is also an expectation of a skew in the site frequency spectrum (SFS) toward both high-and low-frequency derived alleles within the vicinity of the beneficial mutation (Braverman et al. 1995; Simonsen et al. 1995; Fay and Wu 2000). The theoretical expectations under this model of a single, recent selective sweep have been well described (e.g., Kim and Stephan 2002; Kim and Nielsen 2004), and a composite likelihood ratio (CLR) test was developed by Kim and Stephan (2002) based on these expectations that detects such local reductions in nucleotide diversity and skew in the SFS along the chromosome. This signature is used to identify candidate loci that have experienced the recent action of positive selection by comparing the probability of the observed SFS under the standard neutral model with that under the model of a selective sweep. Subsequent work demonstrated that certain demographic histories may be problematic however, with, for example, severe population bottlenecks often replicating patterns of positive selection (Jensen et al. 2005). Thus, to help reduce this issue, Nielsen et al. (2005) adapted the CLR method for genome-wide data utilizing a null model instead derived from the empirically observed SFS, which they termed SweepFinder (along with a more recent implementation, SweepFinder2; DeGiorgio et al. 2016).

Unlike signatures of selective sweeps, those of balancing selection (see the reviews of Fijarczyk and Babik 2015; Bitarello et al. 2023) – a term that encapsulates a variety of selective processes that maintain genetic variability in populations – can potentially last for considerable timescales (Lewontin 1987). Indeed, the temporal history of a balanced allele has been split into multiple phases, with detectable genomic signatures varying in each phase. For example, Fijarczyk and Babik (2015) characterized these phases as recent (<0.4*N_e_* generations), intermediate (0.4-4*N_e_* generations), and ancient (>4*N_e_* generations), where *N_e_* is the effective population size. The initial trajectory of a newly introduced mutation under balancing selection is indistinguishable from that of a partial selective sweep (Soni and Jensen 2024a), whereby the newly arisen mutation rapidly increases to its balanced frequency, conditional on escaping stochastic loss. The signatures of these partial sweeps include a potential excess of intermediate frequency alleles, extended linkage disequilibrium (LD) owing to the associated genetic hitchhiking effects (see the reviews of Crisci et al. 2013; Charlesworth and Jensen 2021), and weaker genetic structure at genes experiencing balancing selection (Schierup et al. 2000). Once the balanced frequency is reached, the allele under balancing selection fluctuates about this frequency, with recombination breaking up the aforementioned LD patterns (Wiuf et al. 2004; Charlesworth 2006; Pavlidis et al. 2012). If balancing selection persists in species with a divergence time predating the expected coalescent time, the allele under selection may continue to segregate as a trans-species polymorphism (Klein et al. 1998; Leffler et al. 2013). To facilitate detection of long-term balancing selection, Cheng and DeGiorgio (2020) developed a class of CLR-based methods for detecting balancing selection that utilizes a mixture model, combining the expectation of the SFS under neutrality with the expectation under balancing selection, to infer the expected SFS at both a putatively selected site and at increasing distances away from that site, released under the BalLeRMix software package (Cheng and DeGiorgio 2020). As with SweepFinder2, this class of methods utilizes a null model directly derived from the empirical SFS in an attempt to account for deviations from the standard neutral expectation in a model-free manner. Such approaches have been shown to be well-powered in detecting long-term balancing selection (>25*N_e_* generations in age; Soni and Jensen 2024a), depending, as they do, on new mutations accruing on the balanced haplotype and thereby generating the expected skew in the SFS toward intermediate frequency alleles.

### Inferring positive and balancing selection in non-human primates

As one might expect, the majority of scans for positive selection in primates have focused upon human population genomic data. However, a number of studies have found signals of putative positive selection in non-human primates, chiefly in the great apes (The Chimpanzee Sequencing and Analysis Consortium 2005; Enard et al. 2010; Locke et al. 2011; Prüfer et al. 2012; Scally et al. 2012; Bataillon et al. 2015; McManus et al. 2015; Cagan et al. 2016; Munch et al. 2016; Nam et al. 2017; Schmidt et al. 2019), as well as in biomedically-relevant species such as rhesus macaques (The Rhesus Macaque Genome Sequencing and Analysis Consortium et al. 2007) and vervet monkeys (Pfeifer 2017b), often with contradictory results, likely owing to differing methodological approaches as well as to high false-positive rates related to a neglect of demographic effects. For example, Enard et al. (2010) proposed a variant of the Hudson-Kreitman-Aguadé (HKA) test (Hudson et al. 1987) and applied it to chimpanzee, orangutan, and macaque population genomic data, finding a high number of orthologous genes exhibiting simultaneous signatures of selective sweeps; conversely, Cagan et al. (2016) utilized a variety of test statistics for detecting positive selection across differing time scales, finding relatively little overlap between species.

Although the first whole-genome, short-read assembly for the aye-aye was published over a decade ago (Perry et al. 2012), and a more recent long-read assembly was made available in 2023 (Shao et al. 2023), the lack of protein-coding gene annotations has greatly limited the ability to scan for recent, episodic selective events in the species. However, the release of a fully annotated, chromosome-level hybrid *de novo* assembly (based on a combination of Oxford Nanopore Technologies long-reads and Illumina short-reads, and scaffolded using genome-wide chromatin interaction data) earlier this year (Versoza and Pfeifer 2024) provides a unique opportunity to identify patterns of episodic selection in this endangered species. In this study, we have performed scans for selective sweeps and balancing selection using SweepFinder2 (DeGiorgio et al. 2016) and BalLeRMix (Cheng and DeGiorgio 2020), respectively, utilizing unique, high-quality, whole-genome, population-level data. Importantly, to account for the confounding effects of demography (Barton 1998; Ewing and Jensen 2014; Poh et al. 2014; Harris and Jensen 2020; and see Charlesworth and Jensen 2024), we used the demographic model of Terbot et al. (2024) which was generated using non-functional regions in the aye-aye genome that are free from the effects of background selection. This demographic history fits neutral population data exceedingly well, and suggests a history in which the aye-aye population size was greatly reduced with first human contact in Madagascar 3,000 -5,000 years ago, with an additional decline owing to recent habitat loss over the past few decades. This well-fitting null model is thus utilized here to determine thresholds for positive and balancing selection scans across functional regions in order to avoid extreme false-positive rates (Thornton and Jensen 2007; Poh et al. 2014; Johri et al. 2022a,b, 2023; Soni et al. 2023) – a particularly important feature in this application given the severe bottleneck history of the species (Terbot et al. 2024). Through this process, we have identified a number of candidate loci with evidence of positive and balancing selection effects, and discuss these results in light of the recently available gene annotations.

## RESULTS & DISCUSSION

We sequenced the genomes of five unrelated aye-aye (*D. madagascariensis*) individuals (three females and two males) housed at the Duke Lemur Center to an average coverage of >50x. After mapping reads to the recently published chromosome-level genome assembly for the species, we called variant and invariant sites following the best practices for non-model organisms (Pfeifer 2017a; van der Auwera and O’Connor 2020). We ran genomic scans across this newly generated population-level dataset, using the 14 autosomal scaffolds from the aye-aye genome assembly of Versoza and Pfeifer (2024) (excluding the X-chromosome, i.e., scaffold 9). The CLR methods implemented in SweepFinder2 and the *B_0MAF_* statistic of the BalLeRMix software package were used to infer selective sweeps and balancing selection, respectively. Selective sweep inference was performed at each single nucleotide polymorphism (SNP), whilst balancing selection inference was performed in windows of size 10 and 100 SNPs. Figures 1 and 2 provide the results of the genome-wide scans for SweepFinder2 and *B_0MAF_* based on 100 SNP windows, respectively (and see Supplementary Figure S1 for the genome-wide scan results with *B_0MAF_* on 10 SNP windows), highlighting a number of peaks along the likelihood surface for each analysis.

**Figure 1:**
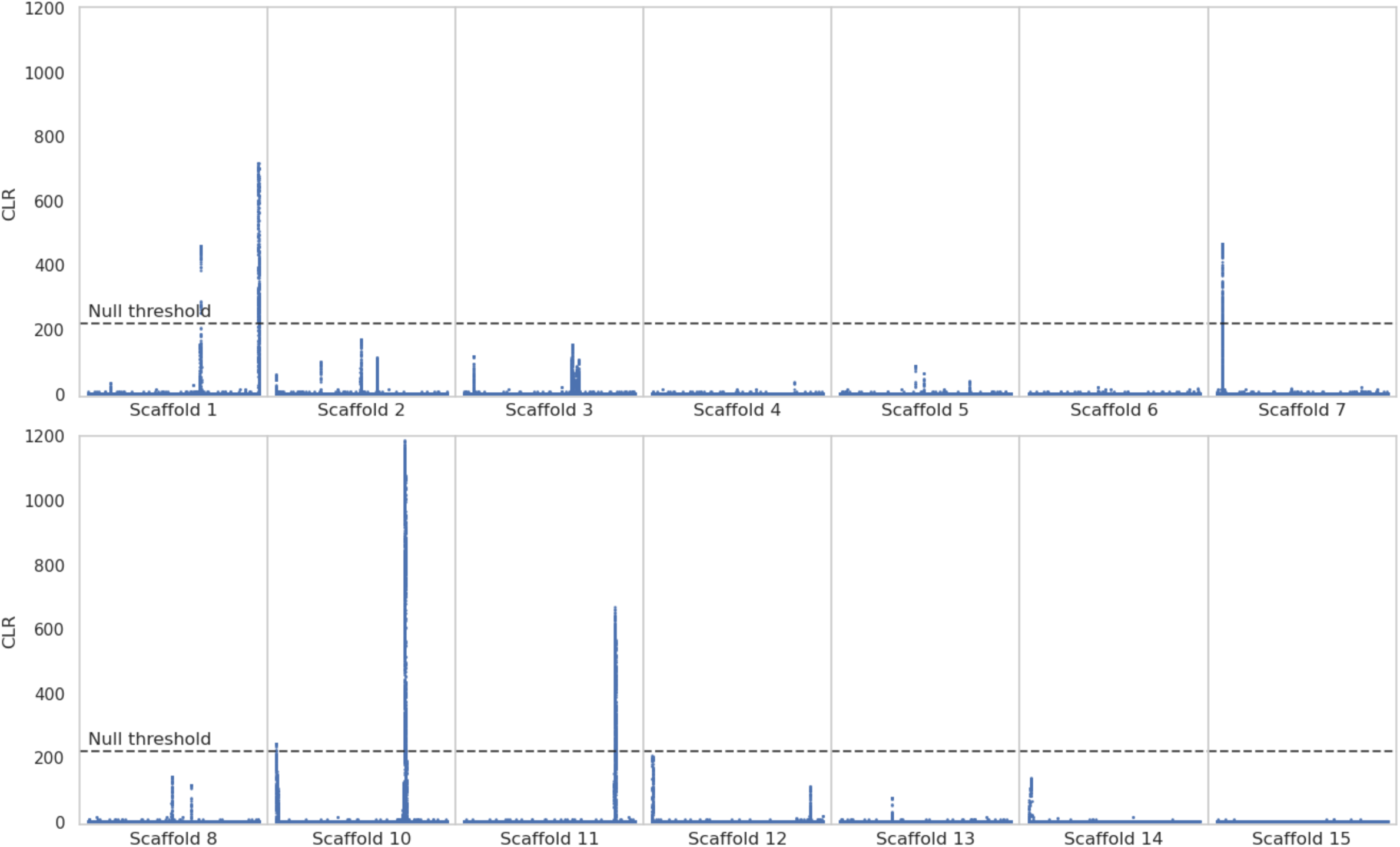
Genome scans for selective sweeps using SweepFinder2. Blue data points are CLR values inferred at each SNP. The dashed line is the threshold for sweep detection, determined by the highest CLR value across 100 simulated replicates of each of the 14 autosomal scaffolds (see "Materials and Methods" section for further details). The x-axis represents the position along the scaffold, and the y-axis represents the CLR value at each SNP.

**Figure 2:**
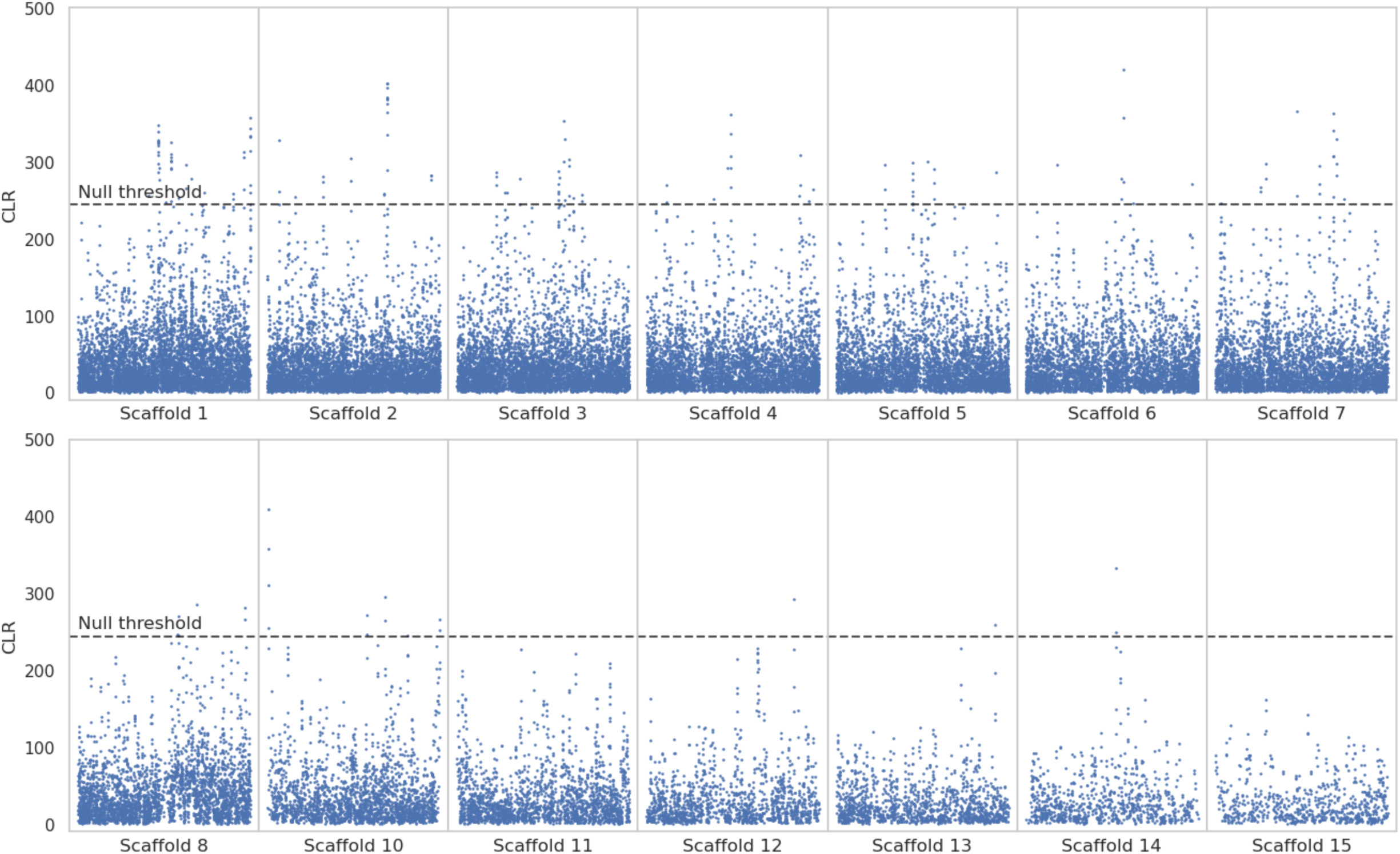
Genome scans for balancing selection using the *B_0MAF_* method. Blue data points are CLR values inferred over windows of length 100 SNPs. The dashed line is the threshold for detection, determined by the highest CLR value across 100 simulated replicates of each of the 14 autosomal scaffolds (see "Materials and Methods" section for further details). The x-axis represents the position along the scaffold, and the y-axis represents the CLR value at each window.

Although it is common to use outlier approaches to identify candidate regions experiencing positive selection, it has previously been shown that such approaches are often associated with extreme false-positive rates (Teshima et al. 2006; Thornton and Jensen 2007; Jensen et al. 2008; Jensen 2023; Soni et al. 2023). Furthermore, these approaches are problematic as any evolutionary model (including standard neutrality) will naturally have a 1% and 5% tail, and thus assuming that genes in these tails of an observed empirical distribution are likely sweep candidates is inherently flawed (see Harris et al. 2018). Moreover, it is not a given that recent sweeps, if they exist, will necessarily even appear in the tails of the empirical distribution under any given demographic model. Thus, we instead followed the recent recommendations of Johri et al. (2022b) in carefully constructing an evolutionarily appropriate baseline model accounting for commonly acting evolutionary processes, including a well-fit population history based on neutral genomic data, purifying and background selection acting on functional sites as modelled via a realistic distribution of fitness effects (DFE), as well as underlying mutation and recombination rate heterogeneity. Furthermore, we characterized the expected true-and false-positive rates of the utilized statistical approaches for our given genomic dataset, as positive and balancing selection will not necessarily be detectable within the context of any given baseline model (Barton 1998; Thornton and Jensen 2007; Poh et al. 2014; Harris and Jensen 2020); importantly, even if these events are not detectable, this approach remains necessary for managing false-positive rates.

We therefore simulated 100 replicates for each of the 14 autosomes using msprime (Baumdicker et al. 2022), under the aye-aye demographic model inferred by Terbot et al. (2024), utilizing the genome assembly of Versoza and Pfeifer (2024), and modelling the specific data details of our newly presented whole-genome dataset. We performed SweepFinder2 and *B_0MAF_* analyses on these baseline simulations, using the maximum CLR values across all simulation replicates as the null thresholds for positive and balancing selection inference, reasoning that these are the maximum values that can be generated in the absence of episodic selective processes under the baseline model considered, and any CLR values that exceeded the thresholds in our empirical analyses were considered to represent meaningful candidate regions. The identified threshold values under this model were 211.747 for SweepFinder2 inference at each SNP, 50.817 for *B_0MAF_* inference on 10 SNP windows, and 244.382 for *B_0MAF_* inference on 100 SNP windows.

### Signatures of episodic selection in the aye-aye genome

A total of 3,462 loci met our null threshold for selective sweep inference using SweepFinder2, which mapped to 71 genes within the aye-aye genome. Scaffolds 1, 7, 10, and 11 contained at least one candidate region, although numerous regions on other scaffolds show peaks that were below our null threshold. For balancing selection inference with *B_0MAF_*, no windows met our null threshold for windows of size 10 SNPs, and 163 windows met our null threshold for windows of size 100 SNPs. The latter windows mapped to 60 candidate genes, covering all autosomal scaffolds apart from scaffolds 11, 14, and 15. Supplementary File S1 provides tables of candidate regions overlapping genes exhibiting CLR values greater than the null thresholds for these analyses.

We manually curated the 71 sweep candidate genes, particularly noting those associated with the peak of each significant likelihood surface (e.g., see Figure 3 for scaffolds 1 and 7 for zoomed inset plots, and Supplementary Figures S2-S3 for additional scaffolds containing candidate regions). Because balancing selection candidate genes were based on 100 SNP windows, these peaks were already quite localized by comparison, though we similarly manually curated this associated set of 60 candidate genes (see Figure 4 for the results in scaffold 1 with mapped genes marked on the plot, and Supplementary Figures S4-S12 for additional scaffolds containing candidate regions). The gene that exhibited the strongest signal of positive selection (i.e., the highest CLR value) was SMPD4 on scaffold 10, whilst the gene exhibiting the strongest signal of balancing selection was LRP1B on scaffold 6 for the 100 SNP window analysis. Biallelic loss-of-function variants in SMPD4 have been found to cause microcephaly, a rare and severe neurodevelopmental disorder with progressive congenital microcephaly and early death in humans (Magini et al. 2019; Smits et al. 2023). LRP1B is a putative tumor suppressor (Brown et al. 2021), and one of the most altered genes in human cancer (Principe et al. 2021). Mutations in LRP1B have been associated with an increased tumor mutation burden (Yu et al. 2022). In one of the earliest large-scale primate genome scans, Nielsen et al. (2005) found that a number of genes involved in tumor suppression were identified as positive selection candidates in humans, as well as genes involved in spermatogenesis, which our analysis in aye-ayes also identified (see below).

**Figure 3:**
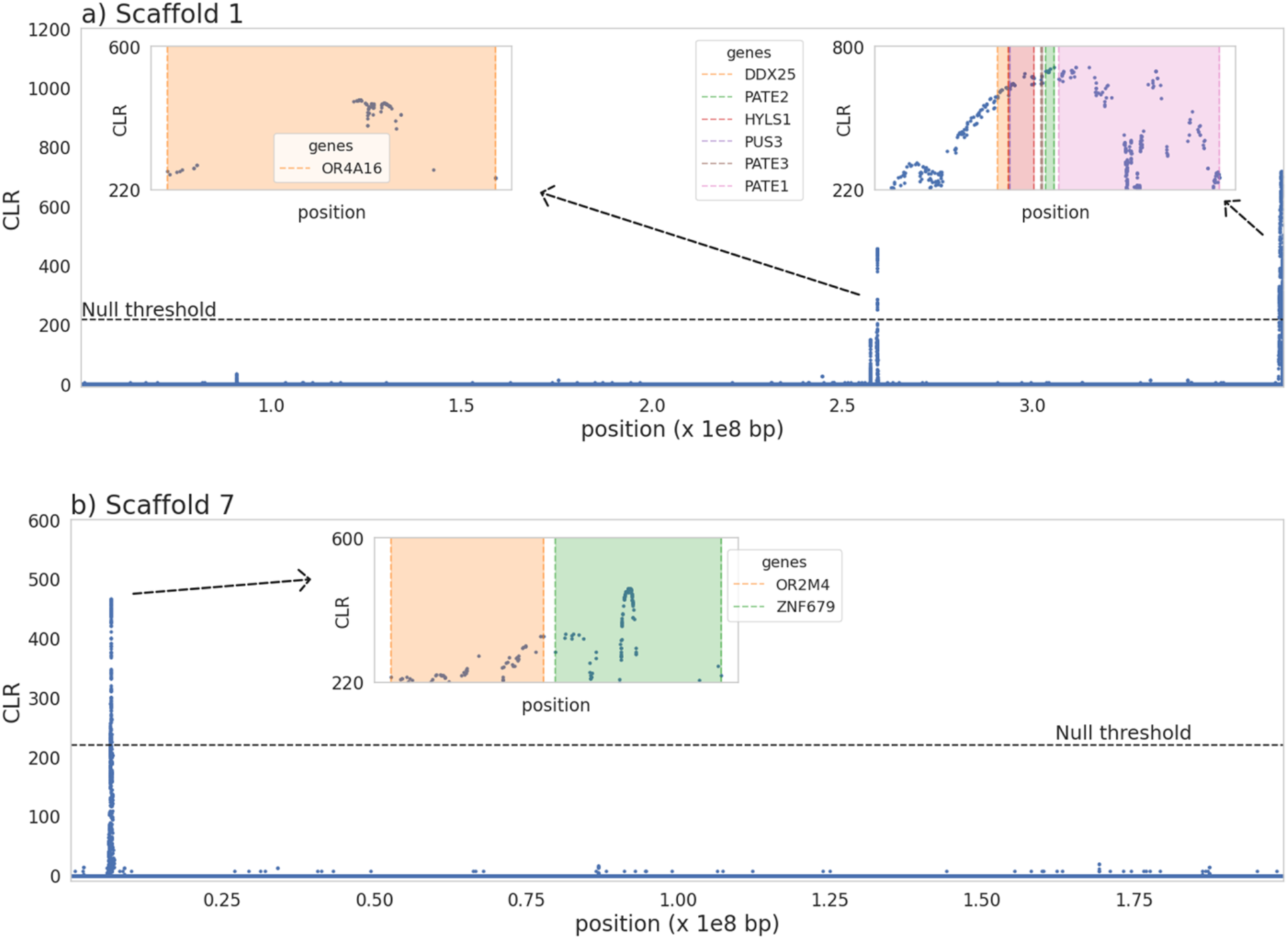
SweepFinder2 selective sweep scan results for a) scaffold 1 and b) scaffold 7. Inset plots zoom in on likelihood surface peaks, with genes in these regions highlighted. The x-axis represents the position along the scaffold, and the y-axis represents the CLR value at each SNP.

**Figure 4:**
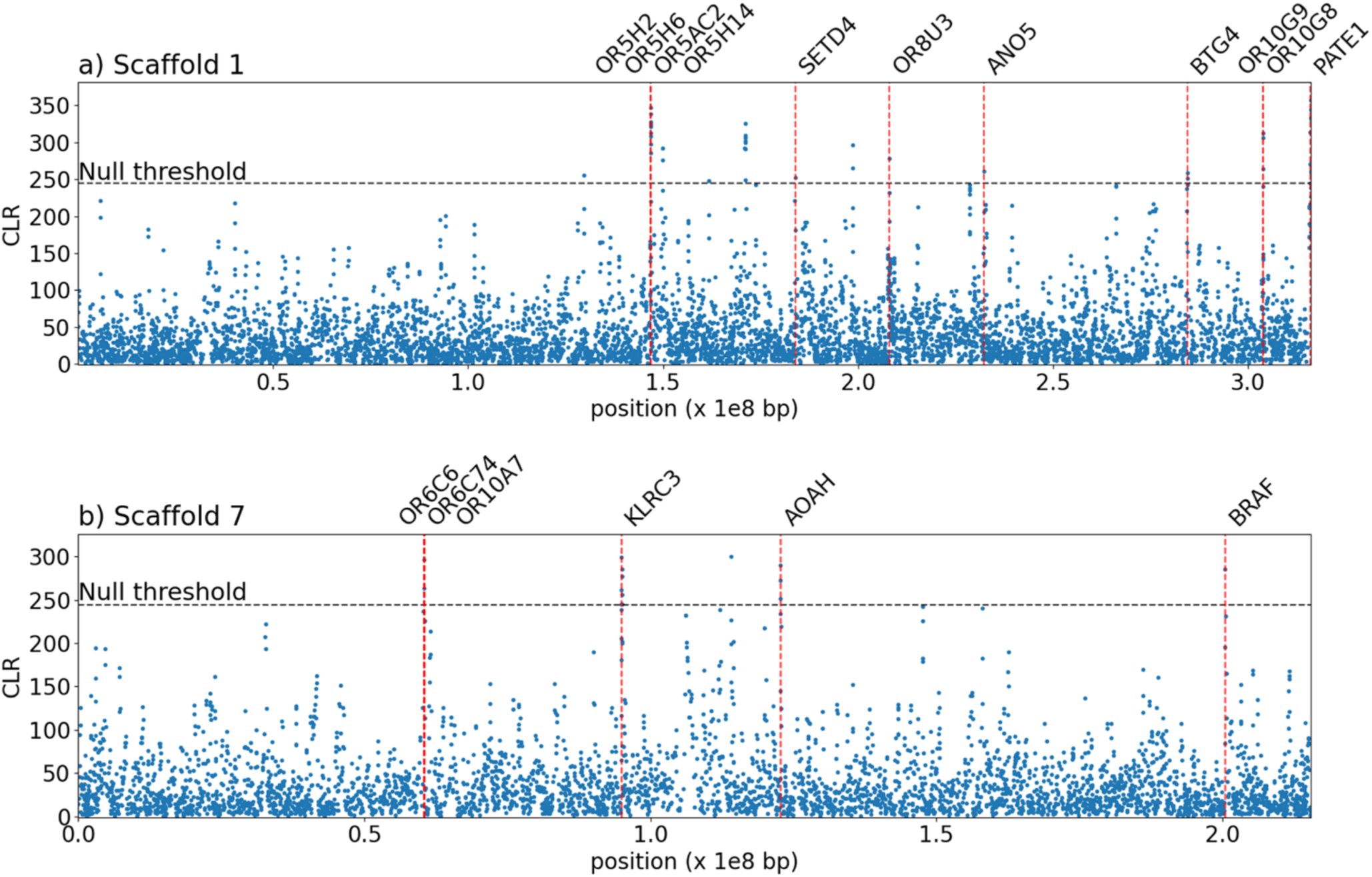
*B_0MAF_* balancing selection scan results for 100 SNP window analysis on a) scaffold 1 and b) scaffold 7. Red vertical lines map to candidate genes. Instances where CLR values meet the null threshold, but no gene is denoted, indicates that no gene overlap was found. The x-axis represents the position along the scaffold, and the y-axis represents the CLR value of each 100 SNP window.

A single candidate region (scaffold 1:315,975,047-315,981,365) in the 100 SNP window balancing selection analysis was found to overlap with a 89,250bp inversion (scaffold 1:315,972,759-316,062,009). This region may represent a false positive, owing to the reduced recombination related to the inversion potentially generating long haplotype structure (Stevison et al. 2011). However, inversions may themselves be selectively maintained, particularly if they contain a beneficial combination of alleles (e.g., Hager et al. 2022; and see Villoutreix et al. 2021); thus, this candidate region will require future dissection.

### Gene functional analysis

A gene function analysis using the Database for Annotation, Visualization, and Integrated Discovery (DAVID; Ma et al. 2023) predicted an enrichment for 19 Gene Ontology (GO) terms in aye-ayes at a *p* ≤ 0.05 for recent selective sweep candidate genes – however, no terms passed a false discovery rate (FDR) threshold of 0.05 or 0.1 for selective sweep candidate genes. Conversely, 14 of the 43 GO terms identified for the 100 SNP window analysis passed both the *p* ≤ 0.05 and the FDR ≤ 0.05 thresholds for balancing selection. Table 1 provides the top 12 functional categories for balancing selection candidate genes for the 100 SNP analysis (and see Supplementary File S1 for all enriched categories). These categories show considerable enrichment and the majority are involved in olfaction.

**Table 1:**
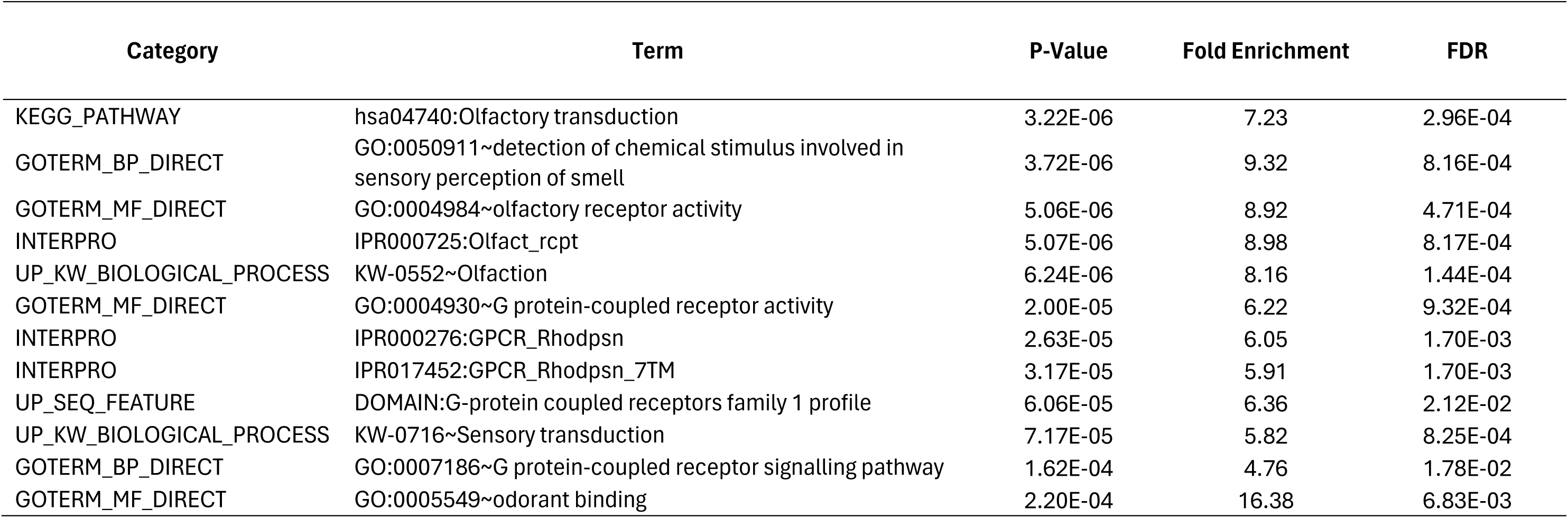
Top 12 hits from gene functional analysis with DAVID on balancing selection candidate genes from 100 SNP window analysis.

### Olfaction

Seven enriched gene functions for balancing selection (enrichment score >1) were related to olfaction, all with high-fold enrichment (ranging from 5.82 to 16.38). A number of OR genes were also found to meet our null threshold for selective sweeps. OR genes provide the basis for the sense of smell (e.g., Buck and Axel 1991), and comprise the largest gene superfamily in mammalian genomes (Glusman et al. 2001; Zozulya et al. 2001). Although olfaction plays a role in locating food, mating, and avoiding danger, the importance and sensitivity of smell varies significantly even amongst closely related species, and it has been suggested that this gene family is subject to a birth-and-death model of evolution, whereby new genes are formed by gene duplication and some of the duplicate genes differentiate in function, whilst others become inactive or are removed from the genome (Niimura and Nei 2003, 2005a,b, 2007). Indeed, OR genes within primates show relaxed selective constraints in apes relative to Old and New World monkeys (Dong et al. 2009), and whilst patterns of variability in chimpanzees are consistent with purifying selection acting on intact OR genes, patterns in humans have been suggested to indicate the action of positive selection (Gilad et al. 2003a). These findings were partially corroborated by Williamson et al. (2007), who found evidence of recent selective sweeps in numerous OR genes in human populations. Gilad et al. (2003a) argued that the differing selective processes acting on OR genes in humans and chimpanzees are likely reflections of differences in lifestyles between humans and other great apes, resulting in distinct sensory needs. Further studies have suggested that both positive selection (Gilad et al. 2003b) and balancing selection (Alonso et al. 2008) are acting on OR genes in humans. Importantly, it has been proposed that heterozygosity in ORs can increase the number of different odorant-binding sites in the genome (Lancet 1994), and thus heterozygote advantage may be an important process. Given that aye-ayes have been shown to discriminate based on scent (Price and Feistner 1994) and use scent-marking behaviors to attract mates (Winn 1994), OR genes represent interesting candidate loci for having experienced on-going positive and balancing selection.

### Rhodopsin

Six enriched gene functions for balancing selection (enrichment score >1) were related to G protein coupled receptors (GPCR), with fold enrichment scores ranging from 3.75 to 6.22. GPCRs are cell surface receptors responsible for detecting extracellular molecules and activating cellular responses; consequently, they are involved in numerous physiological processes. Two of the six enriched GPCR functions were specifically related to rhodopsin, which is the opsin responsible for mediating dim light vision (Litman and Mitchell 1996). It has previously been shown that, despite being nocturnal, aye-ayes maintain dichromacy – potentially supporting previous work that found that the red/green opsin gene survived the long nocturnal phase of mammalian evolution, which has been hypothesized to relate to a role in setting biorhythms (Nei et al. 1997). These identified candidate regions may also lend credence to the speculation that dichromatic nocturnal primates may be able to perceive color while foraging under moonlight conditions (Perry et al. 2007).

### PATE gene family

The PATE gene family has been shown to express in the testis, encoding a sperm-related protein (Bera et al. 2002). Of the four genes that make up the PATE gene family, three were found in candidate selective sweep regions in our analysis (Figure 3), whilst PATE1 was also found in our balancing selection scans based on 100 SNP windows (Figure 4). Soler-Garcia et al. (2005) found that PATE is highly expressed in the male genital tract, and that proteins are secreted into the semen, suggesting a potential in mammalian sperm maturation, whilst Margalit et al. (2012) found that PATE proteins are involved in sperm-oolemma fusion and penetration.

Although primate sperm displays a general uniformity, previous studies have found variations in sperm morphology (Cummins and Woodall 1985; Gage 1998), often predicated on the absence or presence of sperm competition. Indeed, the use of coagulated ejaculate that forms sperm plugs to avoid sperm competition and increase male fertilization success has been described in multiple species of primates, particularly those exhibiting polygynandrous mating systems (i.e., multi-male, multi-female mating systems; Dixson et al. 2002; Martinez and Garcia 2020). As aye-ayes are polygynandrous (Quinn and Wilson 2004), these candidate loci may similarly be hypothesized to relate to mate competition / sexual selection shaping the PATE family of genes.

### Zinc-finger genes

Multiple zinc-finger (ZNF) genes were identified as having undergone recent selective sweeps or balancing selection in our scans of the aye-aye genome. ZNF genes are the largest family of transcription factors in mammalian genomes and play an important role in gene regulation. Previous studies have identified a number of KRAB-ZNF genes – a sub-family of the deeply conserved *Kruppel-*type zinc-finger (KZNF) genes (Bellefroid et al. 1993; Looman et al. 2002; Huntley et al. 2006) – with evidence of positive selection in humans (Nielsen et al. 2005; Nowick et al. 2010, 2011; Jovanovic et al. 2021); additional research has suggested that KRAB-ZNF genes follow a species-specific – as opposed to tissue-specific – pattern of expression (Kapopoulou et al. 2016), suggesting that these genes have different tissue preferences in different species and thus have functionally diversified across the primate lineage (Liu et al. 2014). In addition, numerous gene regulatory factors were identified as putative candidate regions, likely related to the well-described roles of these genes in modifying expression patterns (Berrio et al. 2020; Liu and Robinson-Rechavi 2020; Jovanovic et al. 2021; and see the review of McDonald and Reed 2023).

### Quantifying power to detect recent positive and balancing selection in aye-ayes

It has previously been demonstrated that the statistical power to detect positive selection in any given population will depend on a variety of factors, ranging from the details of the population history, to the amount and configuration of the analyzed data itself (e.g., Johri et al. 2021, 2022b; Soni et al. 2023; Soni and Jensen 2024a; Soni et al. 2024b). In order to quantify the specific power of this analysis to detect candidate regions in aye-ayes, we ran forward-in-time simulations in SLiM (Haller and Messer 2023), utilizing the well-fitting demographic history presented in Terbot et al. (2024). As the underlying DFE in aye-ayes remains uncharacterized, functional regions were simulated using the discrete DFE inferred for humans by Johri et al. (2023) and verified by Soni and Jensen (2024b), in order to provide a reasonable proxy of purifying and background selection effects. Additionally, without access to mutation and recombination maps that are available in heavily studied species, we were unable to directly simulate these effects. However, in order to model the impact of this uncertainly as well as likely rate heterogeneity (Johri et al. 2022a; Soni et al. 2024), we drew rates from a uniform distribution for each 1kb window, such that the mean across each simulation replicate was equal to the mean genomic rate (see the "Materials and Methods" section for more details).

In each simulation replicate, a single beneficial mutation was introduced. For selective sweep models, three different selection regimes were considered, with population-scaled strengths of selection, 2*N_e_s*, of 100, 1,000, and 10,000, where *N_e_* is the ancestral population size and *s* is the strength of selection acting on the beneficial mutation. Five different introduction times of the beneficial mutation were considered, *τ* = 0.1, 0.2, 0.5, 1, and 2, where *τ* is the time before sampling in *N* generations. These values were chosen as there is not expected to be power to detect selective sweeps beyond 4*N* generations, though power generally decays much more rapidly (Kim and Stephan 2002; Przeworski 2002, 2003; Ormond et al. 2016). Only simulation replicates in which the beneficial mutation fixed were retained. For balancing selection, the beneficial mutation was modelled as experiencing negative frequency-dependent selection, and introduced at *τ* = 10*N*, 50*N*, and 75*N* generations, as it has been shown that SFS-based methods have little power to detect recent balancing selection (Soni and Jensen 2024a). Only simulation replicates in which the balanced mutation was still segregating at the time of sampling were retained.

Figures 5 and 6 provide ROC plots for selective sweep inference with SweepFinder2 and *B_0MAF_*, respectively. As shown, the power to detect positive selection in aye-ayes is expected to be reasonably poor in all cases. This is unsurprising, given that aye-ayes have undergone a recent bottleneck, an event which may replicate patterns of variation consistent with a selective sweep (e.g., Barton 1998; Poh et al. 2014; Harris and Jensen 2020), followed by a period of further decline, whilst balancing selection inference is expected to be confounded by the resulting skew in the SFS toward intermediate frequency alleles (Soni and Jensen 2024a). Additionally, identifying the episodic and locus-specific patterns of positive selection is more challenging in populations that have experienced bottlenecks owing to the large genealogical variance generated by this event, leading to widely dispersed test statistics across the genome which can give the illusion of a locus-specific pattern (Thornton and Jensen 2007). While power to detect selective sweeps generally related with strength as expected, this relationship was partially off-set by the fact that at higher strengths of selection the beneficial mutation fixed more rapidly in the population, and thus the time since fixation was longer thereby reducing inference power, as has been previously described analytically (Kim and Stephan 2000). By contrast, power to detect balancing selection increased with time since the introduction of the balanced mutation, as has been previously shown (e.g., Soni and Jensen 2024a). In summary, these power analyses provide a key for interpreting our empirical analysis, in demonstrating that any statistically detectable selective sweep would need be both strong and recent, while any statistically detectable loci experiencing balancing selection would need to be relatively ancient.

**Figure 5:**
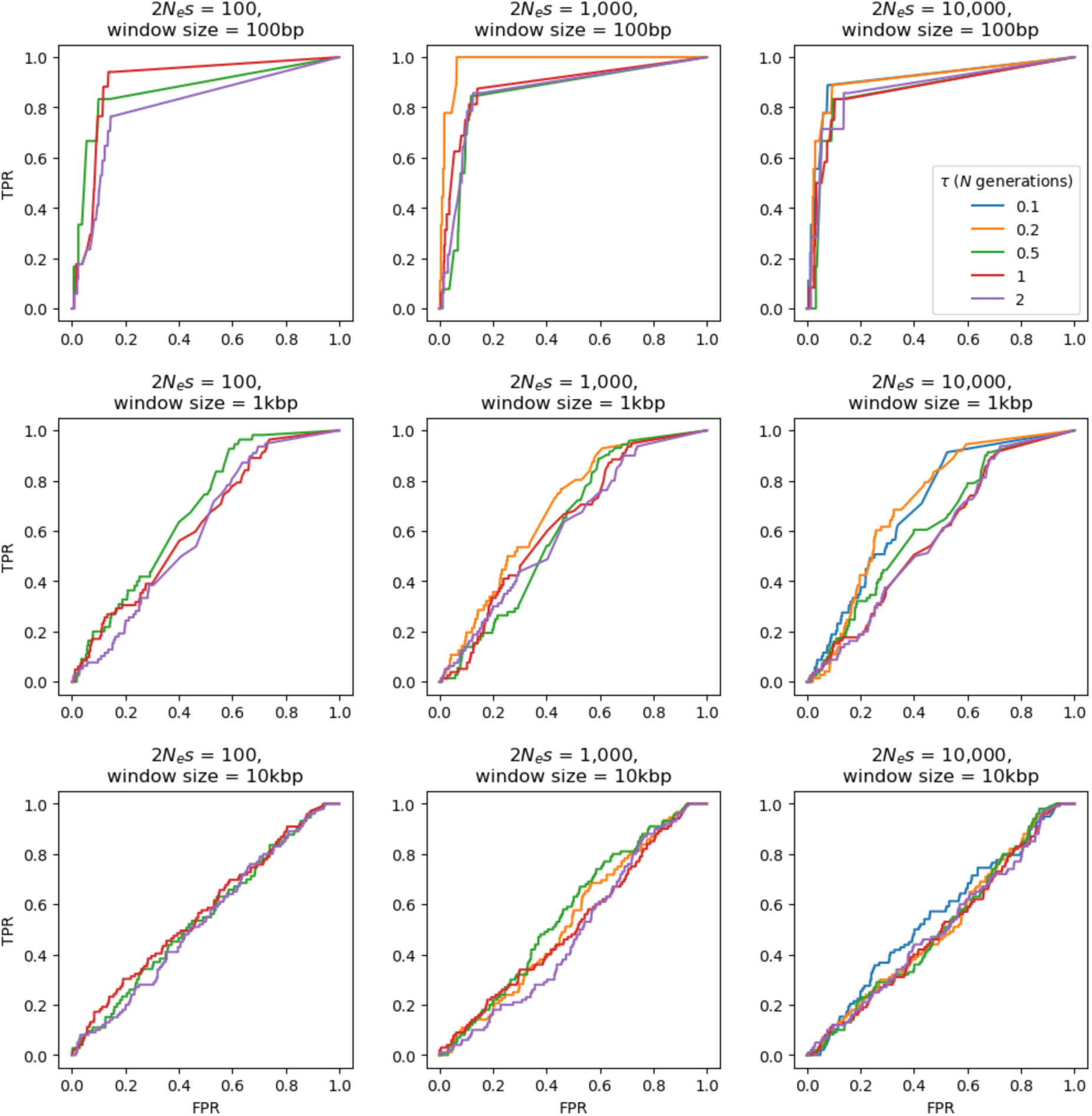
ROC plots for SweepFinder2 showing the change in true-positive rate (TPR) as the false-positive rate (FPR) increases, for sweep inference in aye-ayes across 100 simulation replicates under the Terbot et al. (2024) demographic model, with mutation and recombination rates drawn from a uniform distribution such that the mean rate per simulation rate is equal to the fixed rate (see "Materials and Methods" section). Power analysis was conducted across three selection regimes (population-scaled strengths of selection of 2*N_e_s* = 100, 1,000, and 10,000), five different times of introduction of the beneficial mutation (*τ* = 0.1, 0.2, 0.5, 1, and 2, in *N* generations), and three window sizes (100bp, 1kb, and 10kb). If no ROC is plotted, this is a case in which the beneficial mutation was unable to fix prior to the sampling time in any of the simulation replicates (e.g., at 2*N_e_s* = 100 and *τ* = 0.1).

**Figure 6:**
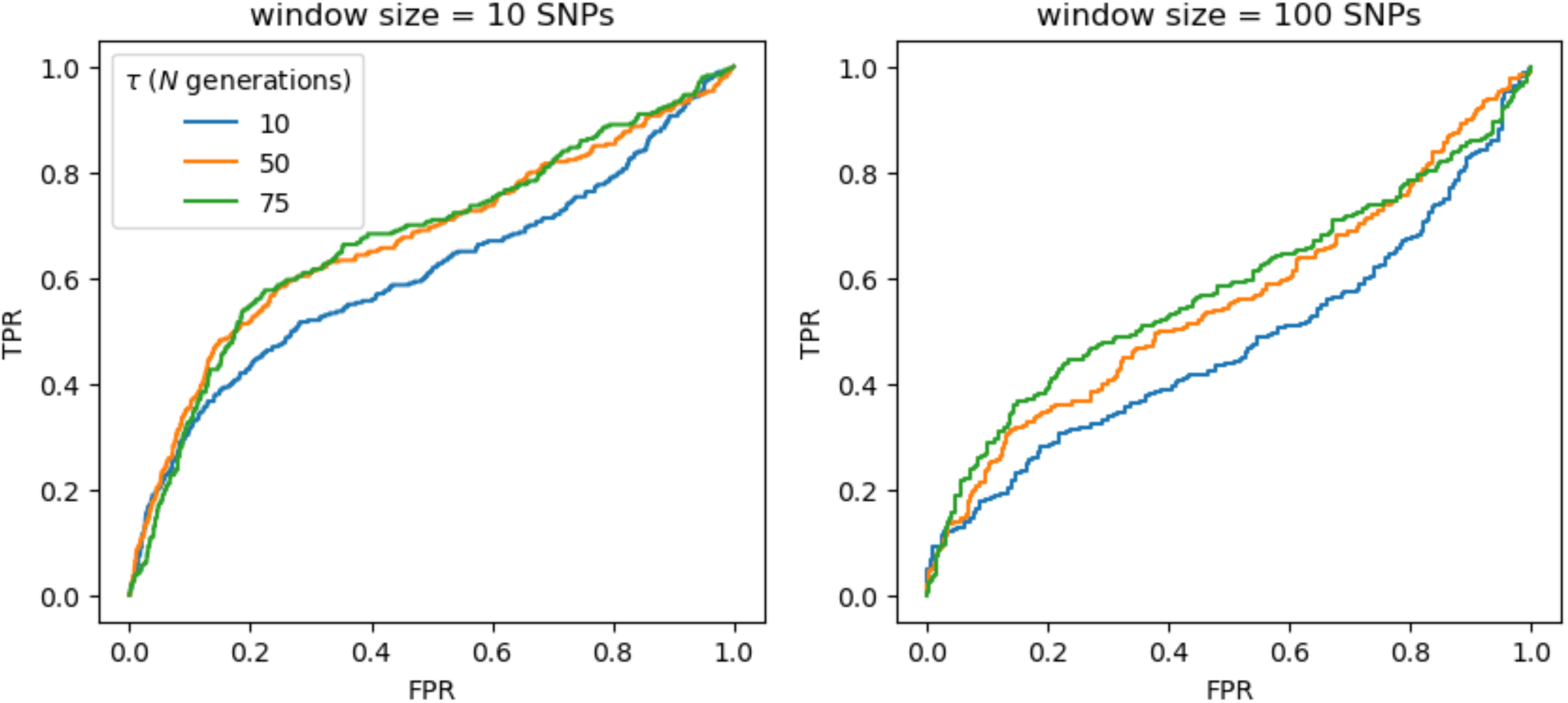
ROC plots for *B_0MAF_* showing the change in true-positive rate (TPR) as the false-positive rate (FPR) increases, for balancing selection inference in aye-ayes across 100 simulation replicates under the Terbot et al. (2024) demographic model, with mutation and recombination rates drawn from a uniform distribution such that the mean rate per simulation rate is equal to the fixed rate (see "Materials and Methods" section). Power analyses were conducted across three different times of introduction of the balanced mutation (*τ* = 10*N*, 50*N*, and 75*N* generations).

### Concluding thoughts

We ran the first large-scale scans for loci having experienced selective sweeps and balancing selection in the aye-aye genome, using newly generated, high-quality, whole-genome, population-level data, and utilizing the recent fully-annotated genome assembly of Versoza and Pfeifer (2024) together with the well-fitting demographic model inferred from non-functional genomic regions of Terbot et al. (2024). By simulating an evolutionarily appropriate baseline model employing these details (Johri et al. 2022a,b), we were able to generate conservative null thresholds for these scans, thereby greatly reducing the types of false-positive rates associated with outlier approaches (Teshima et al. 2006; Thornton and Jensen 2007; Jensen et al. 2008; Soni et al. 2023). The differences between our conservative baseline model approach and a traditional genomic outlier approach are considerable (and see the discussions in Howell et al. 2023; Jensen 2023; Johri et al. 2023; Terbot et al. 2023). For example, selection inference with *B_0MAF_* on 10 SNP windows yielded no candidate windows, and inference on 100 SNP windows yielded 163 candidate windows. An outlier approach interpreting the (commonly used) 5% tail of the empirical distribution of CLR values as candidate loci would instead give 22,971 candidate windows at 10 SNP windows, and 2,280 for 100 SNP windows. More to the point, however, as demonstrated in our power analyses, the great majority of density in these tail distributions may be readily generated by the population history alone, and thus need not invoke positive or balancing selection as an explanation.

Through this more thorough and conservative approach, we have identified a number of promising candidate genes with evidence of having been episodically impacted by positive and balancing selection during the recent evolutionary history of the species, with particularly notable examples being those involved in olfaction and spermatogenesis. Given the on-going destruction of aye-aye habitats, it may well be hypothesized that these vital functions are experiencing changing selection pressures, particularly in light of their solitary lifestyle and polygynandrous mating system.

## MATERIALS AND METHODS

### Animal subjects

This study was approved by the Duke Lemur Center’s Research Committee (protocol BS-3-22-6) and Duke University’s Institutional Animal Care and Use Committee (protocol A216-20-11). The study was performed in compliance with all regulations regarding the care and use of captive primates, including the U.S. National Research Council’s Guide for the Care and Use of Laboratory Animals and the U.S. Public Health Service’s Policy on Human Care and Use of Laboratory Animals.

### Samples and whole-genome sequencing

We sequenced five unrelated aye-aye (*D. madagascariensis*) individuals (three females and two males) using DNA extracted from whole blood samples previously collected at the Duke Lemur Center (Durham, NC, USA). A 150-bp paired-end library was prepared for each sample using the NEBNext Ultra II DNA PCR-free Library Prep Kit (New England Biolabs, Ipswich, MA, USA) and sequenced at high-coverage (>50-fold) on the Illumina NovaSeq platform (Illumina, San Diego, CA, USA) (Supplementary Table S1).

### Calling variant and invariant sites

Calling of variant and invariant sites followed the best practices for non-model organisms as described in Pfeifer (2017a) and van der Auwera and O’Connor (2020). In brief, we pre-processed raw reads by removing adapters and low-quality bases from the read-ends using the Genome Analysis Toolkit (GATK) *MarkIlluminaAdapters* v.4.2.6.1 (van der Auwera and O’Connor 2020) and TrimGalore v.0.6.10 (https://github.com/FelixKrueger/TrimGalore), respectively. We mapped the prepared reads to the aye-aye reference assembly (DMad_hybrid; GenBank accession number: JBFSEQ000000000; Versoza and Pfeifer 2024) using the Burrows Wheeler Aligner (BWA-MEM) v.0.7.17 (Li and Durbin 2009) and marked duplicates using GATK’s *MarkDuplicates* v.4.2.6.1. We further refined the read mappings by performing multiple sequence realignments using GATK’s *RealignerTargetCreator* and *IndelRealigner* v.3.8, recalibrating base quality scores using GATK’s *BaseRecalibrator* and *ApplyBQSR* v.4.2.6.1, and conducting another round of duplicate removal using GATK’s *MarkDuplicates* v.4.2.6.1. As population genomic resources for aye-ayes are limited, no "gold standard" dataset exists for this recalibration step; instead, we utilized a high-confidence training dataset obtained from pedigreed individuals (see Versoza et al. 2024a for details).

We then called variant and invariant sites from high-quality read alignments (’ *--minimum-mapping-quality* 40 ’) using GATK’s *HaplotypeCaller* v.4.2.6.1 with the ’ *pcr_indel_model* ’ parameter set to NONE as a PCR-free protocol was used during library preparation, the ’ *--heterozygosity* ’ parameter set to 0.0005 to reflect species-specific levels of heterozygosity (Perry et al. 2013), and the ’ *-ERC* ’ parameter set to BP_RESOLUTION to output both variant and invariant sites. Next, we jointly assigned genotype likelihoods across all five individuals at all sites (’ *-all-sites* ’) using GATK’s *GenotypeGVCFs* v.4.2.6.1, accounting for species-specific levels of heterozygosity as detailed above. Following the GATK Best Practices, we applied a set of site-level "hard filter" criteria (*i.e.*, QD < 2.0, QUAL < 30.0, SOR > 3.0, FS > 60.0, MQ < 40.0, MQRankSum < -12.5, and ReadPosRankSum < -8.0) to the sites genotyped in all individuals (AN = 34), and applied upper and lower cutoffs on the individual depth of coverage (0.5 × DP_ind_ and 2 × DP_ind_) to remove regions with an unusual read depth indicative of erroneous calls. The resulting dataset was limited to the autosomes and divided into variant (i.e., SNPs) and invariant sites for downstream analyses (Supplementary Table S2).

### Generating null thresholds for selection inference

We simulated the demographic model of Terbot et al. (2024) to generate null thresholds for the inference of selection using the coalescent simulator msprime v1.3.2. (Baumdicker et al. 2022). Although the Terbot et al. (2024) model involves two populations – one population from northern Madagascar, consisting of four individuals previously sequenced at low coverage (Perry et al. 2013), and one population from the rest of the island, consisting of eight individuals previously sequenced at low coverage (Perry et al. 2013) as well as five individuals newly sequenced at high coverage (Terbot et al. 2024) – we here only consider the single population of newly sequenced individuals (*n* = 5), given the much higher data quality of this sample. As such, while the Terbot et al. (2024) work includes only the non-coding regions of these newly sequenced individuals, we here present the full genome data including functional regions as well. The demographic history of this population involves a relatively ancient size reduction (likely associated with human colonization of Madagascar), followed by a period of recent decline up to the current day (likely related to recent habitat loss). Based on this demographic history, we simulated 10 replicates for each of the 14 autosomal chromosomes included in the most recent aye-aye genome assembly (Versoza and Pfeifer 2024), modelling a mutation rate of 1.52e-8 per base pair per generation (i.e., the mutation rate previously reported in another lemur species; Campbell et al. 2021), and a recombination rate of 1e-8 cM/Mb (as recently inferred from pedigree data; Versoza, Lloret-Villas et al. 2024).

To generate null thresholds, we ran both SweepFinder2 v1.0. (DeGiorgio et al. 2016) and the *B_0MAF_* method of Cheng and DeGiorgio (2020) on the allele frequency files generated from our simulated demographic data. In brief, we performed inference at each SNP with SweepFinder2 using the following command: SweepFinder2 –lu GridFile FreqFile SpectFile OutFile. Additionally, we utilized two inference schemes in *B_0MAF_*: (1) windows containing 10 SNPs with a 5 SNP step size and (2) windows containing 100 SNPs with a 50 SNP step size, using the following command: python3 BalLeRMix+_v1.py -I FreqFile --spect SfsFile -o OutFile -w W -s S –usePhysPos –noSub –MAF –rec 1e-8, where *W* is the window size and *S* is the step size, both based on number of SNPs. Because we lacked information on the polarization of SNPs, allele frequencies were folded and only polymorphic sites were included in the analyses. Notably, the highest CLR value across all null model simulations was set as the null threshold for inference, under the assumption that this is the highest value that can be generated in the absence of positive or balancing selection, thereby providing a conservative scan to reduce false positive rates.

### Inferring recent positive and balancing selection in the aye-aye genome

We ran SweepFinder2 and *B_0MAF_* on the 14 aye-aye autosomes using the same inference schema discussed above. Only those inference values greater than the null threshold values were considered as putatively experiencing positive or balancing selection. The genes in which these selected sites were located were identified using the genome annotations of Versoza and Pfeifer (2024), whilst overlaps with structural variants were also identified, based on those described by Versoza et al. (2024b). Because the number of identified genes was relatively small (<200 for both the sweep and balancing selection scans), we manually curated our candidates. In brief, for sweep candidates, we first identified genes under the significant likelihood surface. These candidate genes were then run through the NCBI database (Sayers et al. 2022) and Expression Atlas (Madeira et al. 2022) in order to identify function and expression patterns in other primate species. Additionally, we performed a Gene Ontology analysis (The Gene Ontology Consortium 2023) using the Database for Annotation, Visualization, and Integrated Discovery (Ma et al. 2023) on our candidate genes (conducted for all candidates together, and also separately for selective sweep and balancing selection candidates).

### Power analyses

To assess how much statistical power exists to detect episodic selection in this aye-aye population given the details of both the demographic history and of the dataset itself, we simulated the Terbot et al. (2024) demographic model forward-in-time in SLiM v.4.0.1 (Haller and Messer 2023). Thereby, the simulated region was comprised of three functional regions, separated by intergenic regions of length 16,489bp. Each functional region contained nine exons of length 130bp, separated by introns of length 1,591bp, for a total region length of 91,161bp. These details were estimated from the Versoza and Pfeifer (2024) genome annotations to represent common aye-aye genomic architecture. Mutations in intronic and intergenic regions were modelled as effectively neutral, while exonic mutations were drawn from a DFE comprised of four fixed classes (Johri et al. 2020), whose frequencies are denoted by *fi*: *f0* with 0 ≤ 2*N_e_s* < 1 (i.e., effectively neutral mutations), *f1* with 1 ≤ 2*N_e_s* < 10 (i.e., weakly deleterious mutations), *f2* with 10 ≤ 2*N_e_s* < 100 (i.e., moderately deleterious mutations), and *f3* with 100 ≤ 2*N_e_s* < 2*N_e_* (i.e., strongly deleterious mutations), where *N_e_* is the effective population size and *s* is the reduction in fitness of the mutant homozygote relative to wild-type. Within each bin, *s* was drawn from a uniform distribution. We utilized the DFE inferred from human population genomic data by Johri et al. (2023), and verified in Soni and Jensen (2024b), as a proxy. This modelling of a realistic DFE in functional regions enabled us to account for the effects of purifying and background selection, in addition to population history, when assessing a baseline model of commonly acting evolutionary processes in the species.

To model uncertainty and heterogeneity in the underlying mutation and recombination rate, each 1kb region of the simulated chromosome was assigned a different rate. Rates were drawn from a uniform distribution such that the chromosome-wide average was approximately the fixed rate previously observed in pedigree data (i.e., 1 cM/Mb; Versoza, Lloret-Villas, et al. 2024). For variable recombination rates, the minimum and maximum parameters of the uniform distribution were 0.01 and 10 cM/Mb, respectively (i.e., a 100-fold decrease and a 10-fold increase on the fixed rate). For variable mutation rates the minimum and maximum parameters of the uniform distribution were set at 0.61e-8 and 3.8e-8 (i.e., 0.5x and 2.5x the fixed rate previously reported in lemurs, respectively).

Simulations had a 10*N* generation burn-in time, where *N* is the ancestral population size of 23,706. A further 10*N* generations were then simulated for the sweep analysis, whilst a further 85*N* generations were simulated for the balancing selection analysis. In each simulation replicate, a single positively selected mutation was introduced. For the selective sweep analysis, the beneficial mutation was introduced at *τ* = [0.1, 0.2, 0.5, 1, 2], where *τ* is the time before sampling in *N* generations. Three different beneficial selection coefficients were simulated: 2*N_e_s* = [100, 1,000, 10,000], where *N_e_* is equal to the ancestral population size of 23,706. For the balancing selection analysis, the balanced mutation was introduced at *τ* = [10*N*, 50*N*, 75*N*]. The balanced mutation experienced negative frequency-dependent selection, which was modelled such that the selection coefficient of the balanced mutation was dependent on its frequency in the population: *S_bp_* = *F_eq_* – *F_bp_*, where *S_bp_* is the selection coefficient of the balanced mutation, *F_eq_* is the equilibrium frequency of the balanced mutation (here set to 0.5), and *F_bp_* is the frequency of the balanced mutation in the population. Simulations were structured such that if the selective sweep failed to fix, or the balanced mutation was either fixed or lost from the population, the simulation would restart at the point of introduction of the selected mutation.

Scans for selective sweeps and balancing selection were then performed on the simulated data, as per the procedure discussed above. ROC plots were generated in order to summarize expected performance. Because selective sweep inference was performed on each SNP, 100bp, 1kb and 10kb windows were generated for creating ROC plots for SweepFinder2 results (note that this was not necessary for *B_0MAF_* as SNP-based windows were used for inference).

## ACKNOWLEDGEMENTS

We would like to thank Erin Ehmke, Kay Welser, and the Duke Lemur Center for providing the aye-aye samples used in this study, and members of the Jensen Lab and Pfeifer Lab for helpful discussion. DNA extraction, library preparation, and Illumina sequencing was conducted at Azenta Life Sciences (South Plainfield, NJ, USA). Computations were performed on the Sol supercomputer at Arizona State University (Jennewein et al. 2023) and on the Open Science Grid, which is supported by the National Science Foundation and the U.S. Department of Energy’s Office of Science. This is Duke Lemur Center publication # XXXX.

## FUNDING

This work was supported by the National Institute of General Medical Sciences of the National Institutes of Health under Award Number R35GM151008 to SPP and the National Science Foundation under Award Number DBI-2012668 to the Duke Lemur Center. VS, JT, and JDJ were supported by National Institutes of Health Award Number R35GM139383 to JDJ. CJV was supported by the National Science Foundation CAREER Award DEB-2045343 to SPP. The content is solely the responsibility of the authors and does not necessarily represent the official views of the National Institutes of Health or the National Science Foundation.

## Notes

### Competing Interest Statement

The authors have declared no competing interest.

