## Supplementary Materials for "A whole-genome scan for evidence of recent positive and balancing selection in aye-ayes (*Daubentonia madagascariensis*) utilizing a well-fit evolutionary baseline model"

**SUPPLEMENTARY TABLES**

| | NCBI | coverage |
|---|---|---|
| | DMad_01 | 104.9 |
| | DMad_02 | 50.5 |
| | DMad_03 | 50.2 |
| | DMad_04 | 53.7 |
| | DMad_05 | 52.5 |

**Supplementary Table S1**. Samples and their sequencing coverage.

| scaffold | # SNPs | # invariant sites | Ts/Tv |
|---|---|---|---|
| 1 | 310,744 | 202,589,728 | 2.49 |
| 2 | 282,464 | 187,007,285 | 2.53 |
| 3 | 266,594 | 166,053,610 | 2.46 |
| 4 | 221,453 | 141,877,129 | 2.51 |
| 5 | 213,515 | 139,978,054 | 2.57 |
| 6 | 201,488 | 134,246,862 | 2.63 |
| 7 | 197,047 | 125,387,556 | 2.56 |
| 8 | 160,561 | 106,567,069 | 2.56 |
| 10 | 111,122 | 74,651,313 | 2.66 |
| 11 | 101,805 | 68,490,964 | 2.88 |
| 12 | 68,829 | 42,837,589 | 2.89 |
| 13 | 66,853 | 39,350,940 | 2.84 |
| 14 | 42,375 | 22,947,744 | 2.77 |
| 15 | 36,095 | 19,884,749 | 2.76 |
| ∑ or ∅ | **2,280,945** | **1,471,870,592** | **2.60** |

**Supplementary Table S2**. Summary of the variant and invariant sites. After filtering, a total of >2.2 million autosomal, biallelic, single nucleotide polymorphisms (SNPs) with a transition-transversion ratio (Ts/Tv) of 2.60 were discovered in the accessible genome.

**SUPPLEMENTARY FIGURES**

## S1

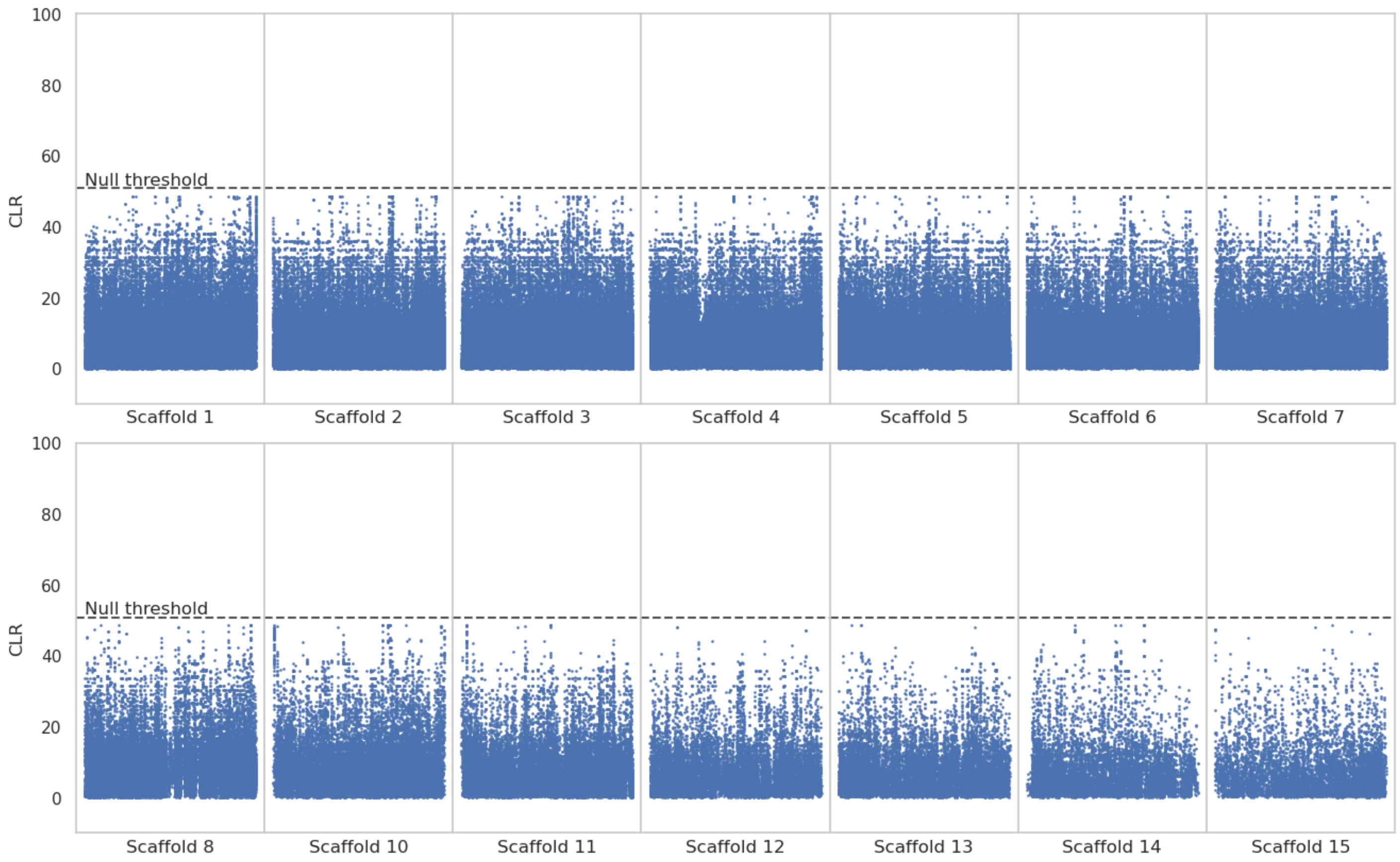


**Supplementary Figure S1:** Genome scans for balancing selection using the $B_{0MAF}$ method. Blue data points are CLR values inferred over windows of length 10 SNPs. The dashed line is the threshold for detection, determined by the highest CLR value across 100 simulated replicates of each of the 14 autosomal scaffolds (see "Materials and Methods" section for further details). The x-axis represents the position along the scaffold, and the y-axis represents the CLR value at each window.

**S2**

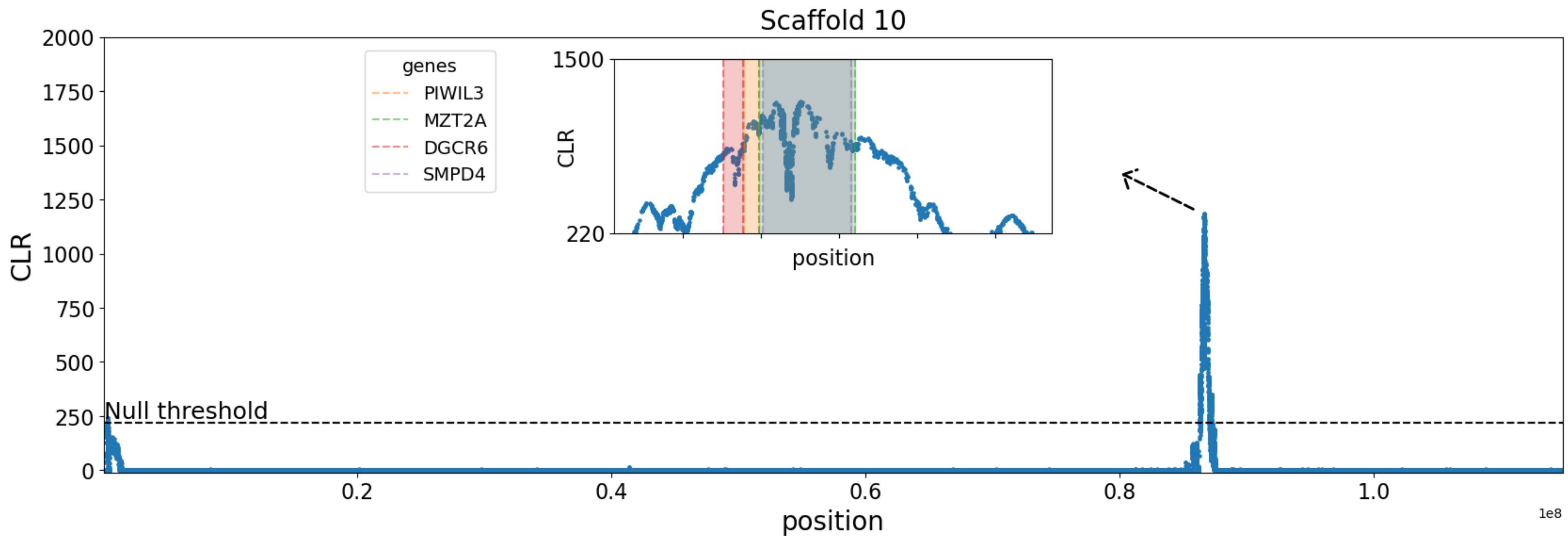


**Supplementary Figure S2:** SweepFinder2 selective sweep scan results for scaffold 10. Inset plot zooms in on likelihood surface peaks, with genes in these regions highlighted. The x-axis represents the position along the scaffold, and the y-axis represents the CLR value at each SNP.

**S3**

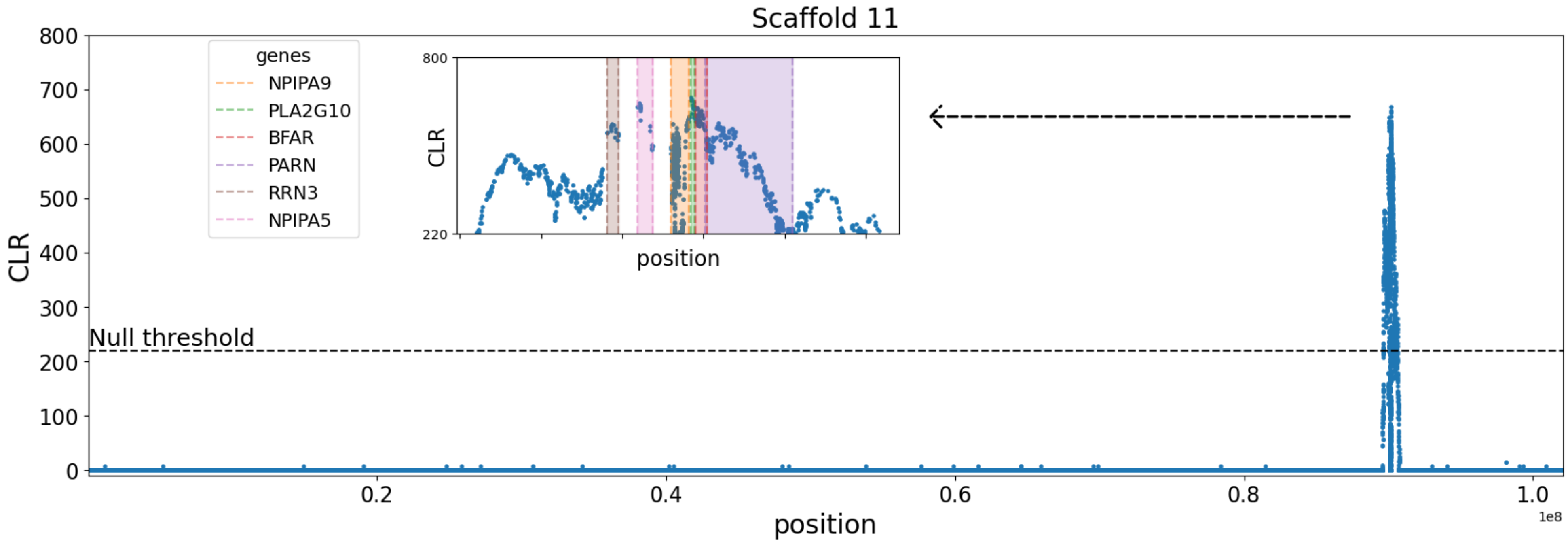


**Supplementary Figure S3:** SweepFinder2 selective sweep scan results for scaffold 11. Inset plot zooms in on likelihood surface peaks, with genes in these regions highlighted. The x-axis represents the position along the scaffold, and the y-axis represents the CLR value at each SNP.

**S4**

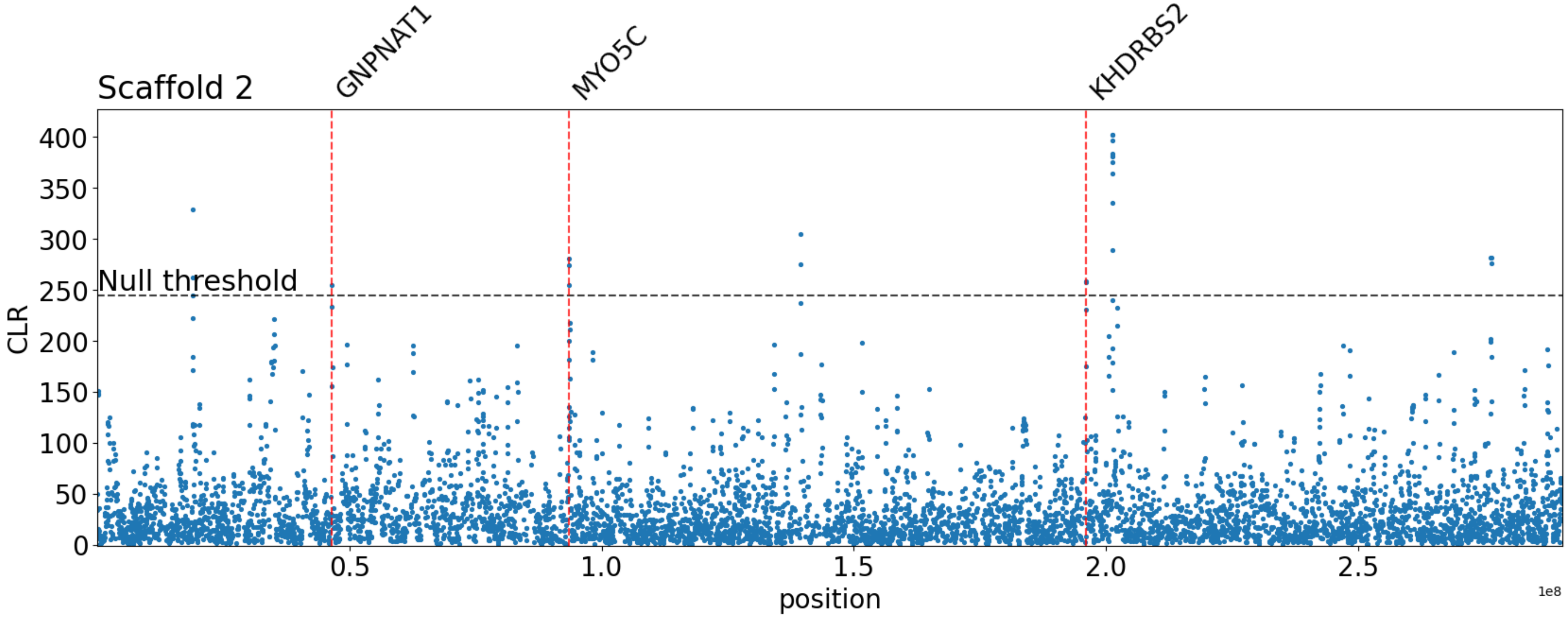


**Supplementary Figure S4:** *$B_{0MAF}$* balancing selection scan results for 100 SNP window analysis on scaffold 2. Red vertical lines map to candidate genes. Instances where CLR values meet the null threshold, but no gene is denoted, indicates that no gene overlap was found. The x-axis represents the position along the scaffold, and the y-axis represents the CLR value of each 100 SNP window.

**S5**

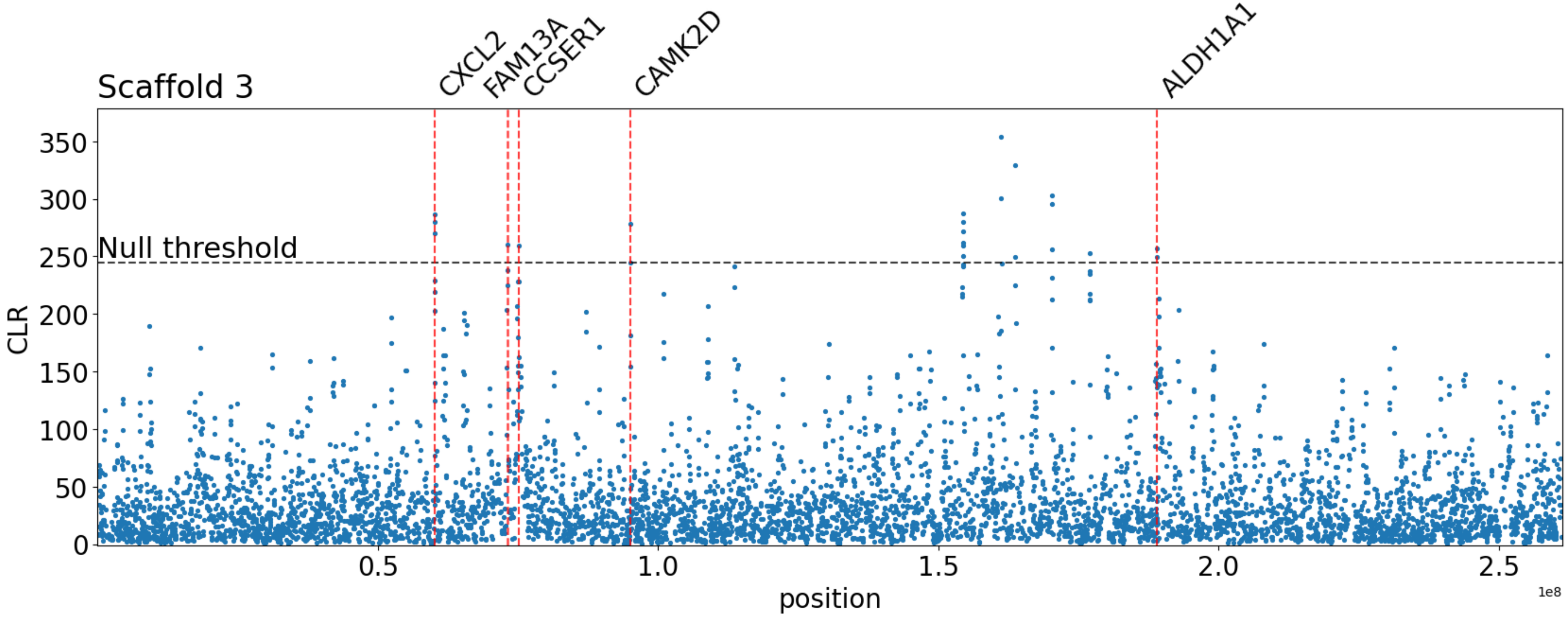


**Supplementary Figure S5:** $B_{0MAF}$ balancing selection scan results for 100 SNP window analysis on scaffold 3. Red vertical lines map to candidate genes. Instances where CLR values meet the null threshold, but no gene is denoted, indicates that no gene overlap was found. The x-axis represents the position along the scaffold, and the y-axis represents the CLR value of each 100 SNP window.

**S6**

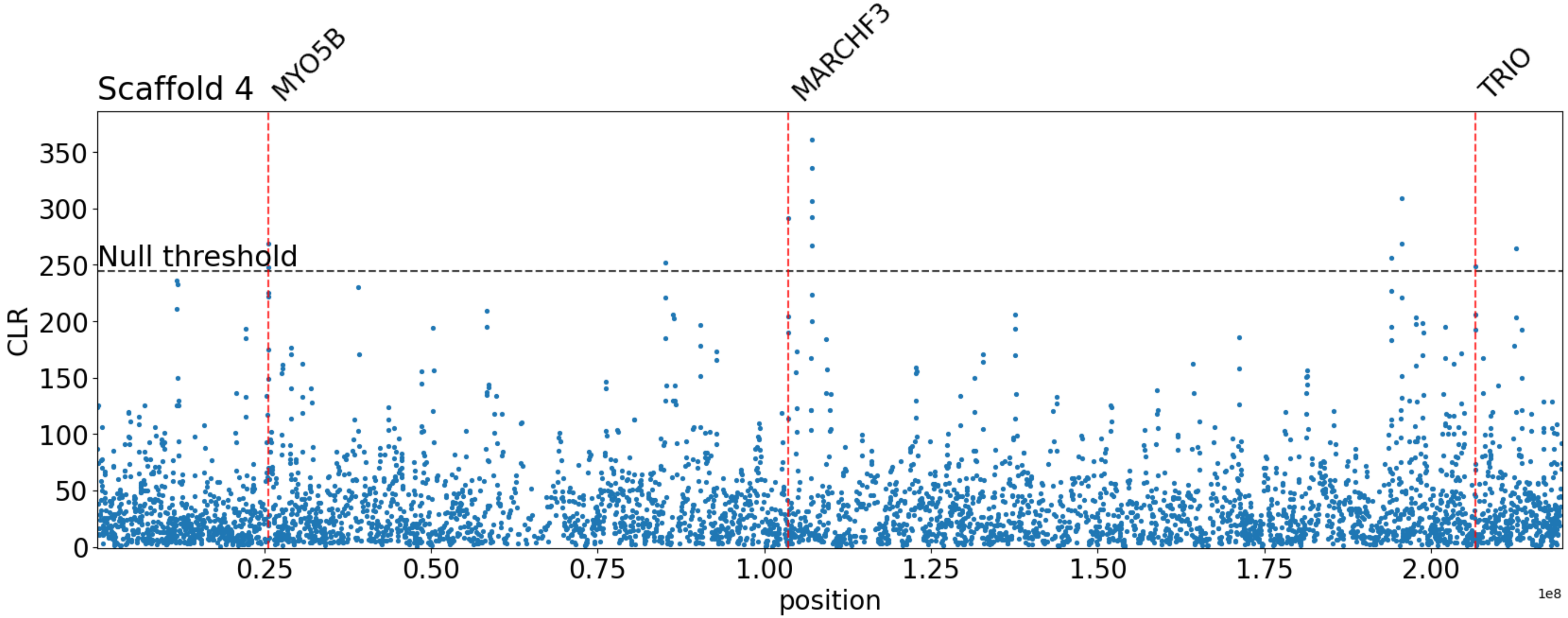


**Supplementary Figure S6:** $B_{0MAF}$ balancing selection scan results for 100 SNP window analysis on scaffold 4. Red vertical lines map to candidate genes. Instances where CLR values meet the null threshold, but no gene is denoted, indicates that no gene overlap was found. The x-axis represents the position along the scaffold, and the y-axis represents the CLR value of each 100 SNP window.

**S7**

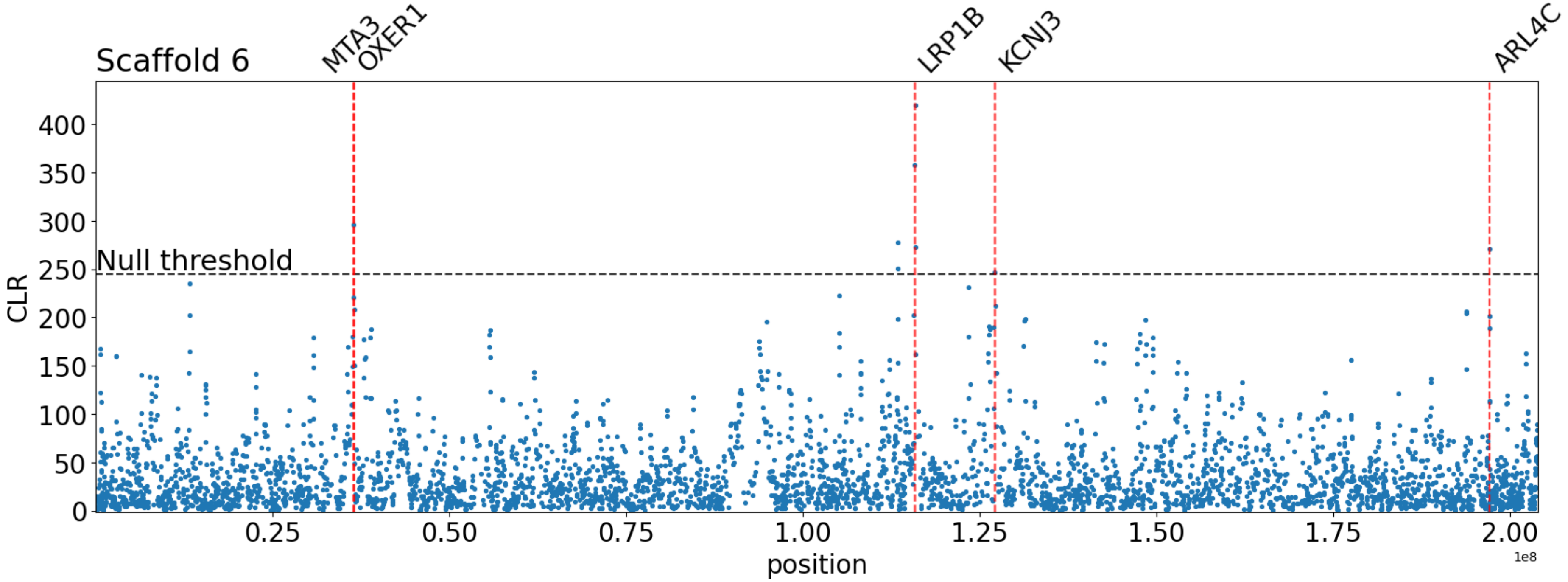


**Supplementary Figure S7:** $B_{0MAF}$ balancing selection scan results for 100 SNP window analysis on scaffold 6. Red vertical lines map to candidate genes. Instances where CLR values meet the null threshold, but no gene is denoted, indicates that no gene overlap was found. The x-axis represents the position along the scaffold, and the y-axis represents the CLR value of each 100 SNP window.

**S8**

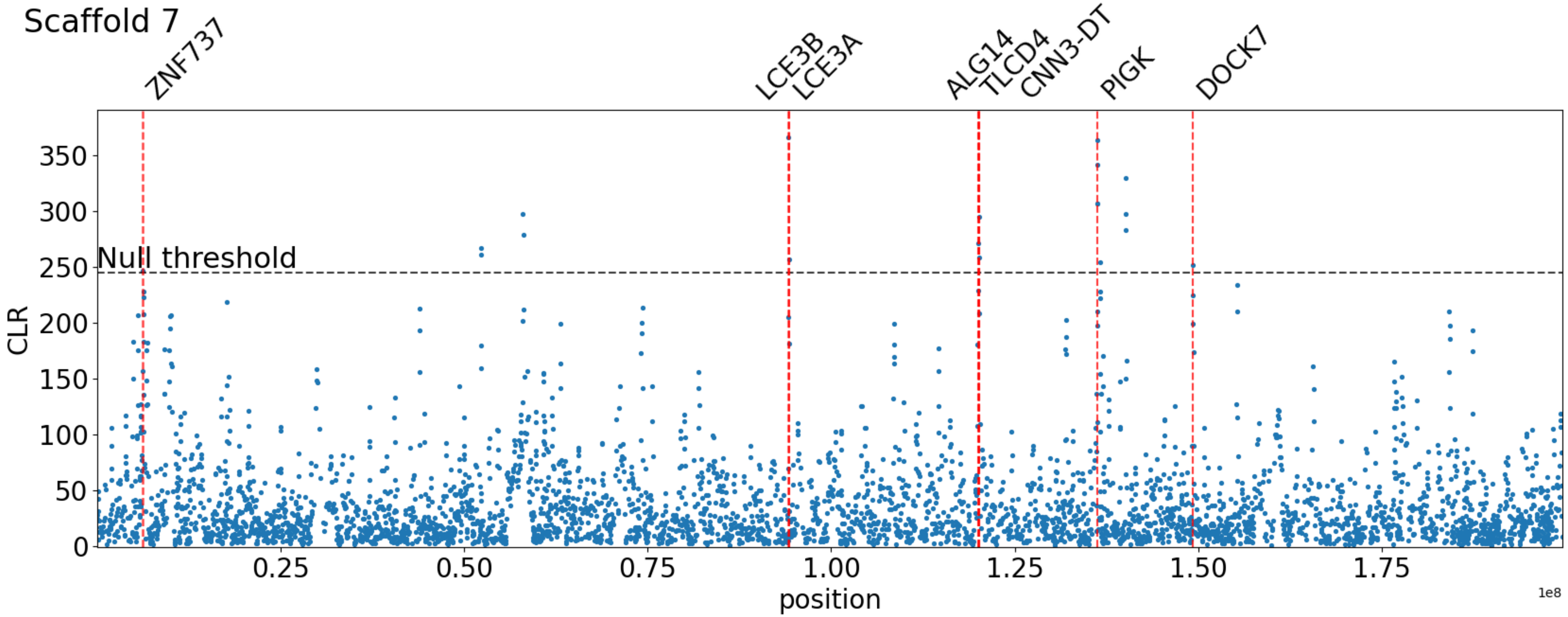


**Supplementary Figure S8:** $B_{0MAF}$ balancing selection scan results for 100 SNP window analysis on scaffold 7. Red vertical lines map to candidate genes. Instances where CLR values meet the null threshold, but no gene is denoted, indicates that no gene overlap was found. The x-axis represents the position along the scaffold, and the y-axis represents the CLR value of each 100 SNP window.

**S9**

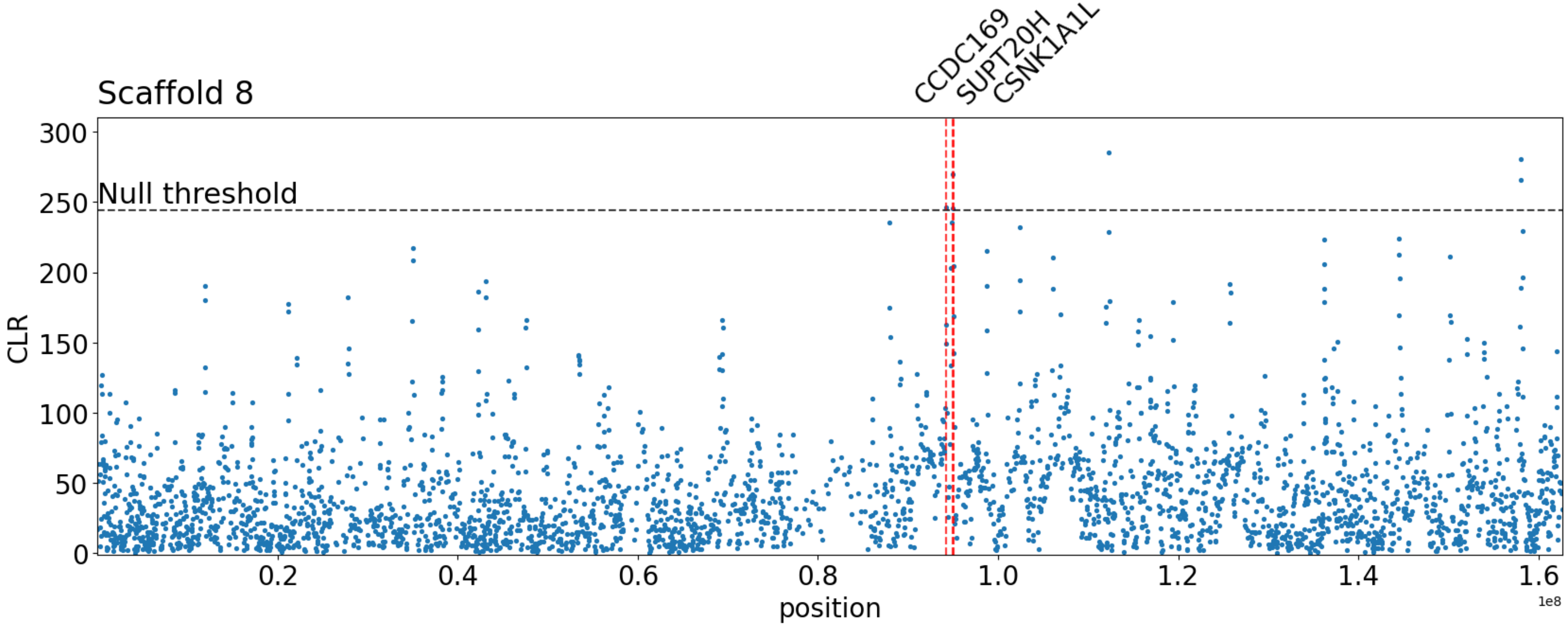


**Supplementary Figure S9:** $B_{0MAF}$ balancing selection scan results for 100 SNP window analysis on scaffold 8. Red vertical lines map to candidate genes. Instances where CLR values meet the null threshold, but no gene is denoted, indicates that no gene overlap was found. The x-axis represents the position along the scaffold, and the y-axis represents the CLR value of each 100 SNP window.

**S10**

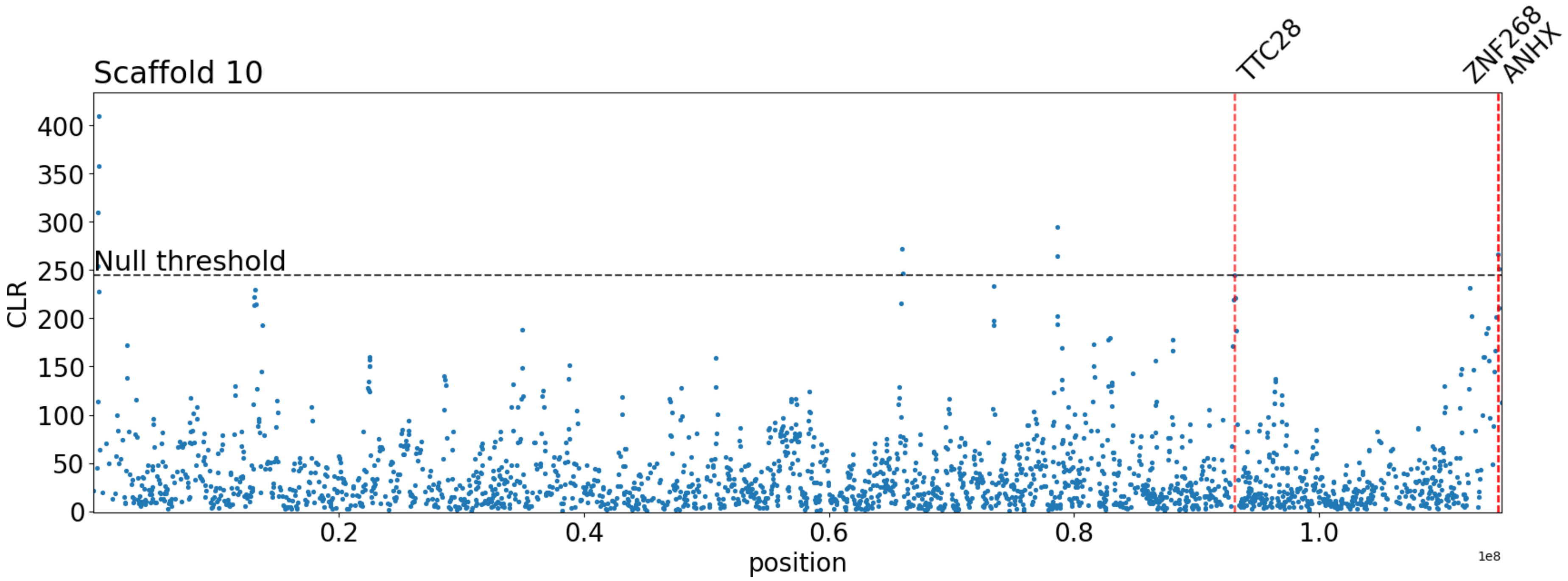


**Supplementary Figure S10:** $B_{0MAF}$ balancing selection scan results for 100 SNP window analysis on scaffold 10. Red vertical lines map to candidate genes. Instances where CLR values meet the null threshold, but no gene is denoted, indicates that no gene overlap was found. The x-axis represents the position along the scaffold, and the y-axis represents the CLR value of each 100 SNP window.

**S11**

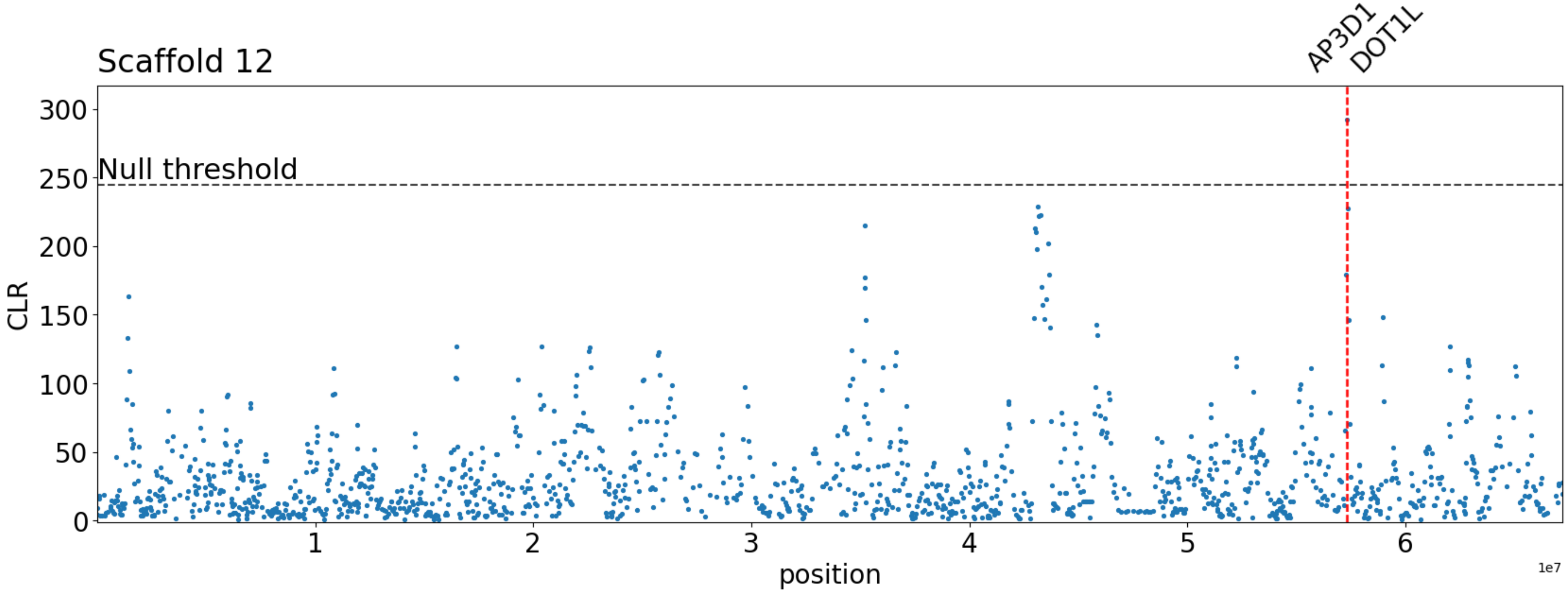


**Supplementary Figure S11:** $B_{0MAF}$ balancing selection scan results for 100 SNP window analysis on scaffold 12. Red vertical lines map to candidate genes. Instances where CLR values meet the null threshold, but no gene is denoted, indicates that no gene overlap was found. The x-axis represents the position along the scaffold, and the y-axis represents the CLR value of each 100 SNP window.

**S12**

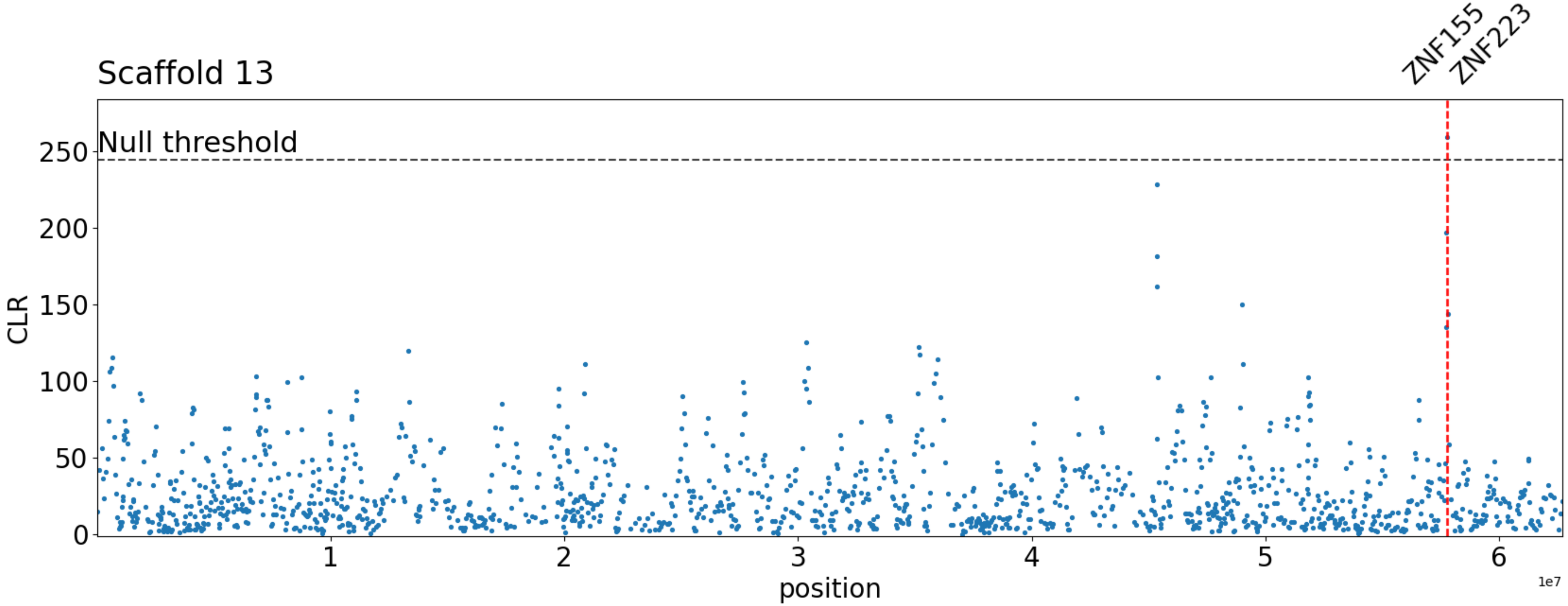


**Supplementary Figure S12:** $B_{0MAF}$ balancing selection scan results for 100 SNP window analysis on scaffold 13. Red vertical lines map to candidate genes. Instances where CLR values meet the null threshold, but no gene is denoted, indicates that no gene overlap was found. The x-axis represents the position along the scaffold, and the y-axis represents the CLR value of each 100 SNP window.
